# Genotype imputation accuracy and the quality metrics of the minor ancestry in multi-ancestry reference panels

**DOI:** 10.1101/2023.05.30.542466

**Authors:** Mingyang Shi, Chizu Tanikawa, Hans Markus Munter, Masato Akiyama, Satoshi Koyama, Kohei Tomizuka, Koichi Matsuda, Gregory Mark Lathrop, Chikashi Terao, Masaru Koido, Yoichiro Kamatani

**Author notes:** Correspondence: M.K. and Y.K. Fax: None.

## Abstract

Large-scale imputation reference panels are now available and have contributed to efficient genome-wide association studies through genotype imputation. However, it is still under debate whether large-size multi-ancestry or small-size population-specific reference panels are the optimal choices for under-represented populations. We imputed genotypes of East Asian (EAS; 180k Japanese) subjects using the Trans-Omics for Precision Medicine (TOPMed) reference panel and found that the standard imputation quality metric (Rsq) substantially overestimated the dosage r^2^ (squared correlation between imputed dosage and true genotype). Variance component analysis of Rsq revealed that the increased imputed-genotype certainty (dosages closer to 0, 1, or 2) caused upward bias, indicating some systemic bias in the imputation. Through systematic simulations using different template switching rates (θ value) in the hidden Markov model, we uncovered that the lower θ value increased the imputed-genotype certainty and Rsq; however, dosage r^2^ was insensitive to the θ value, thereby causing a deviation. In simulated reference panels with different sizes and ancestral diversities, the θ value estimates from Minimac decreased with the size of a single ancestry and increased with the ancestral diversity. Thus, Rsq could overestimate or underestimate dosage r^2^ for a subpopulation in the multi-ancestry panel and the deviation represents different imputed-dosage distributions. Finally, despite the impact of θ value, distant ancestries in the reference panel contributed only a few additional variants passing a predefined Rsq threshold. We conclude that the θ value has a substantial impact on the imputed dosage and the imputation quality metric value.

## Introduction

Genotype imputation is a cost-efficient technique to expand the number of markers in genetic studies. Li and Stephen’s hidden Markov model (HMM) and the pre-phasing-imputation pipeline are employed by the popular tools Minimac, IMPUTE, and BEAGLE, with slightly different implementations but similar accuracy [1–5].

The imputation accuracy (*e.g.*, the squared correlation between imputed dosage and true genotype; dosage r^2^) [6], the scores generated by software without the true genotype (Rsq and INFO) [3,7,8], and the number of variants passing a predetermined Rsq threshold (high-Rsq variants) are widely used quality metrics to evaluate imputation performance [9]. The performance primarily depends on the reference panel size and genetic similarity between the reference panel and target sample [10]. In studies using the International HapMap Project (HapMap) and the 1000 Genomes Project (1KGP) [11,12], pooling multiple ancestries together to maximize the panel size typically improved the imputation performance compared to using only the matched ancestry [8,13,14]. Deelen et al. combined the population-specific whole genome sequencing (WGS) dataset of the Genome of the Netherlands (GoNL) with the 1KGP and showed higher dosage r^2^ than when using either GoNL or 1KGP alone [15]. Huang et al. combined the UK10K with the 1KGP to generate more variants with high INFO [9].

Moreover, larger multi-ancestry reference panels, such as the Haplotype Reference Consortium (HRC) and the Trans-Omics for Precision Medicine (TOPMed) [16,17], greatly improved the imputation accuracy and increased the number of high-Rsq variants in European (EUR), African (AFR), and Admixed American (AMR) [10,18].

However, constructing large reference panels by combining diverse ancestries is not always beneficial. Small-size population-specific panels, such as Norwegian and Estonian panels [19,20], achieve similar performance to the HRC panel in EUR. Moreover, even though the HRC panel includes all 1KGP samples, the imputation accuracy in non-EUR populations can be inferior to that of the 1KGP panel alone [10,21]. Bai et al. found that adding 27 samples from a different ancestry to the Han Chinese (1KGP-CHB, size = 103) resulted in better imputation accuracy than using the 1KGP-CHB or 1KGP panel [22]. In studies of Asian, combining population-specific WGS datasets with the 1KGP had improved the imputation accuracy in some instances [23–25], but not in others [26–28]. So far, the reason for these discrepancies is unclear.

Two sets of parameters are used in the HMM-based imputation algorithms. Transition probability is governed by the template switching rate (θ) between adjacent markers, which models the recombination and relatedness, and emission probability is determined by the error rate (ε) for each marker, which models genotyping error, gene conversion, and recurrent mutation [3,7]. These parameters are crucial to the HMM for calculating matching probabilities between the template and target during the imputation process, but dosage r^2^ is robust to these parameters [3,5]. Many studies have suggested different Rsq or INFO thresholds to achieve a similar dosage r^2^ [29,30], making it difficult to ascertain the relationship between the imputation process, Rsq, and dosage r^2^.

In this study, we imputed the East Asian (EAS) using the TOPMed reference panel (EAS comprised 1.22% of the total samples) and found that Rsq substantially overestimated dosage r^2^. We introduced novel variance component analysis for Rsq and analytically investigated why Rsq was overestimated and characterized the relationship between the template switching rate used in the HMM, quality metrics, and imputed dosage. Further, we evaluated the θ value, Rsq, dosage r^2^, the deviation between them, and the number of high-Rsq variants from matched or distant ancestry in cases where the target ancestry was the major or minor component of the multi-ancestry panels.

## Material and methods

### Subjects, genotyping, and quality control (QC)

Cohort specifications of Biobank Japan (BBJ-180k), genotyping, and QC are provided in **Supplementary Method 1**. For imputation using the TOPMed panel, we lifted the coordinates of the genotyping array from hg19 to hg38 using LiftOver [31]. Palindromic variants (reference/alternative (ref/alt) alleles were G/C or A/T) and variants not in the TOPMed freeze5b (https://bravo.sph.umich.edu/freeze5/hg38/) were excluded. The ref/alt alleles were swapped and/or reverse-complemented according to the hg38 reference sequence. A total of 515,587 autosomal variants remained.

### Pre-phasing, reference panel construction, and imputation

EAGLE v2.4.1 was used for pre-phasing without an external reference [32]. The 1KGP reference panel was downloaded from https://genome.sph.umich.edu/wiki/Minimac3. Methods used to construct the population-specific panels (named the BBJ1k and JEWEL3k) are provided in **Supplementary Method 2**. We used an in-house server to perform imputation using the 1KGP, BBJ1k, and JEWEL3k panels. Minimac3 v2.0.3 was used to estimate the HMM parameters and prepare m3vcf files.[3] Minimac4 v1.0.2 was used for imputation [33]. Imputation using the TOPMed and HRC panel were performed using the TOPMed and Michigan imputation servers [3,17], respectively.

### Evaluation of the imputation performance using different reference panels

Rsq from the Minimac4’s info file was extracted for the subsequent analyses. Variants with Rsq < 0.3 were removed [34]. Imputation performance was empirically evaluated using 993 samples with WGS (named WGS_993_). QC and processing are elaborated in **Supplementary Method 3**. Coverage of a reference panel was the fraction of variants in WGS_993_ that could be imputed with Rsq ≥ 0.3. Dosage r^2^ was the squared Pearson correlation coefficient between the imputed dosage and true genotype (encoded as 0, 1, or 2). Minor allele frequency (MAF) and minor allele count (MAC) of WGS_993_ were used to stratify variants into bins.

### Quantification of the deviation between Rsq and dosage r^2^

We modeled the relationship between the imputed allelic dosage (𝒚 = (𝑦_*i*_, …, 𝑦_*n*_), 𝑦_#_ ∈ [0,1]) and the true allele (𝒙 = (𝑥_*i*_, …, 𝑥_*n*_), 𝑥_#_ ∈ {0,1}) using the simple linear regression formula:

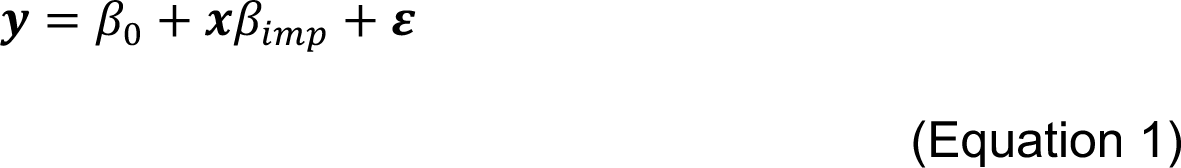

where *β_imp_* is a scalar of the regression coefficient with 𝛽^_#%&_ = 𝐶𝑜𝑣 (𝒙, 𝒚)⁄𝑉𝑎𝑟 (𝒙), *β_0_* is the intercept, and 𝜺 = (𝜖_1_, …, 𝜖_*n*_) is an error term. The Minimac’s EmpRsq metric, which is the squared Pearson correlation coefficient between ***x*** and ***y*** (https://genome.sph.umich.edu/wiki/Minimac3_Info_File), equals the ratio between the regression sum of squares (*SS_reg_*) and the total sum of squares (*SS_tot_*) in this simple linear regression. Then we have:

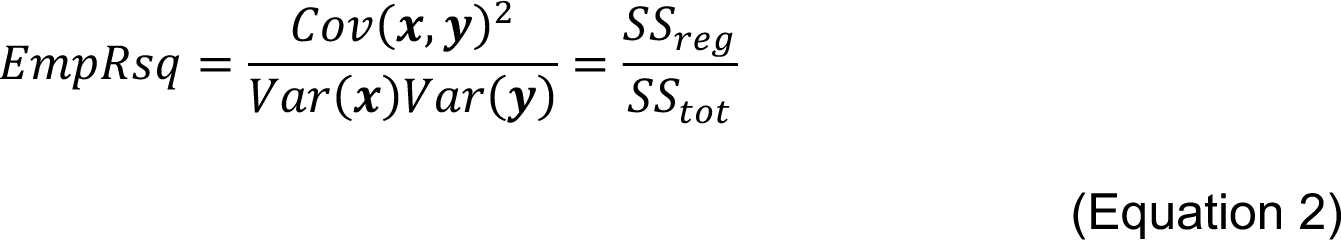

where Cov(***x***, ***y***), Var(***x***), and Var(***y***) are the covariance between ***x*** and ***y***, the variance of ***x***, and the variance of ***y***, respectively.

Rsq is the ratio between Var(***y***) and 𝑝(1 − 𝑝), where *p* is the alternative allele frequency (AAF) in the imputed dataset [3,7]. Then,

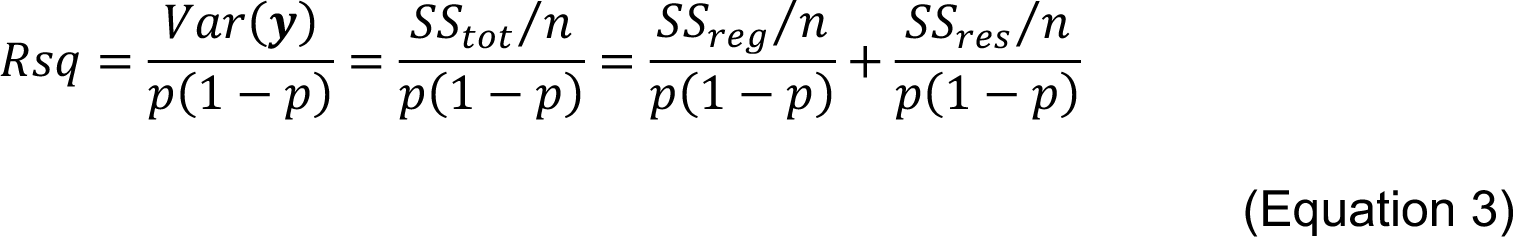

where *SS_res_* is the residual sum of squares that follows *SS_tot_* = *SS_reg_* + *SS_res_*, and *n* is the number of imputed haplotypes. Hence, Rsq comprises two parts: regression-related and residual-related. We define 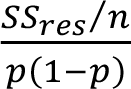 as <u>M</u>AF-<u>A</u>djusted-<u>R</u>esidual-<u>E</u>rror (MARE). Then:

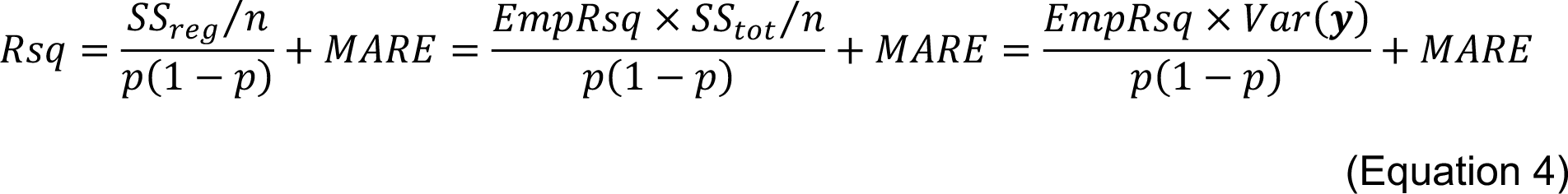

By assuming the equal AAF in ***x*** and ***y***, *i.e.*, 𝑉𝑎𝑟(𝒙) = 𝑝(1 − 𝑝), Rsq could be further treated as:

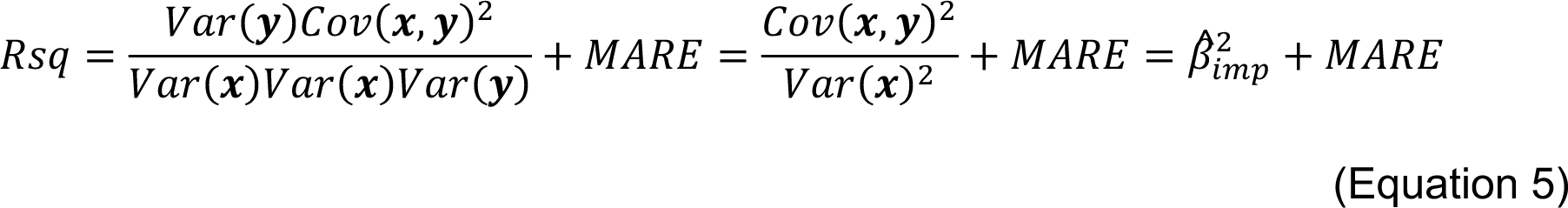

Finally, *MARE* and 𝛽^_#%&_ could be obtained from Rsq and EmpRsq:

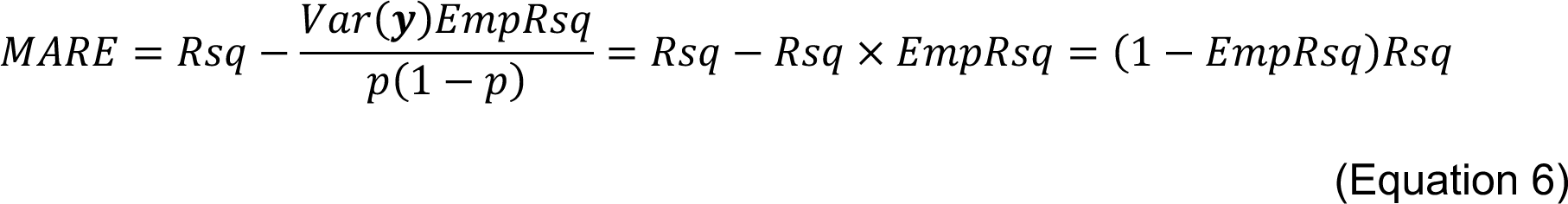

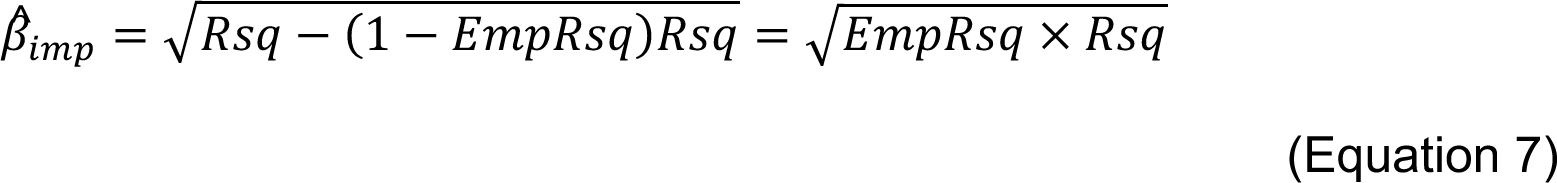

In Equation 7, 𝛽^_#%&_ is negative if ***x*** and ***y*** are negatively correlated. We do not consider that situation. Hence, each combination of Rsq and EmpRsq indicates specific values of *MARE* and 𝛽^_#%&_. Hereafter, we refer to 𝛽^_#%&_ as the β_imp_ metric and *MARE* as the MARE metric.

Because the WGS dataset is unphased diploid data, dosage r^2^ calculated from that will be slightly different from EmpRsq. **Supplementary Note 1** provides the detailed methods to calculate MARE and β_imp_ from haploid or diploid data, and the concordance between values obtained using Equations 6–7 and calculated from the imputed dosage in both haploid and diploid cases.

### Imputation using the simulated 1KGP reference panels with different θ values

The 1KGP vcf files were downloaded from the Minimac3 website (see above). Parameters were estimated using Minimac3. The θ value was extracted (denoted as “Recom” in the m3vcf file) [3], scaled by 21 folds manually (0.01–100), and replaced in the original file. A brief explanation of the θ value is provided in **Supplementary Note 2**. The array data of WGS_993_, a subset of BBJ-180k, was used as the target sample. The imputation procedure was the same as described above.

### Evaluation of the imputation performance using the simulated reference panels

We obtained the imputed allelic dosage (LooDosage) from Minimac4’s empiricalDose file (by turning the “--meta” option on) [35]. It was derived from the leave-one-out (LOO) method by hiding the markers on the array during the imputation. Rsq, EmpRsq, MARE, and β_imp_ were calculated from the LooDosage and array data. MAF was obtained from the array data.

### Simulation of reference panels and estimating the θ value

We sampled subsets from the 1KGP and JEWEL3k, shuffled the sample order 10 times, and created new vcf files using bcftools v1.14 (https://samtools.github.io/bcftools/) [36]. We estimated the parameters using Minimac3. **Supplementary Method 4** provides the detailed methods. The total θ value along chr19 (by summing the θ values between adjacent markers) was used to evaluate the reference panel size and ancestral diversity’s impact. A discussion of the θ value qualification is provided in **Supplementary Note 3**.

### EUR-EAS reference panel simulation and imputation

We randomly sampled 403 individuals (named 1KGP-EUR_403_) from the 1KGP-EUR and combined them with the 1KGP-EAS and 6 subsets (size = 500, 1,000, 1,500, 2,000, 2,500, and 3,256) of the 3,256 JPT WGS samples (named as JPT_3256_) in JEWEL3k. The remaining 100 individuals in the 1KGP-EUR were used as the target sample. We extracted the 10,375 polymorphic variants (chr19) on the Illumina Global Screening Array v3.0 (https://support.illumina.com/content/dam/illumina-support/documents/downloads/productfiles/global-screening-array-24/v3-0/infinium-global-screening-array-24-v3-0-a1-b151-rsids.zip) to simulate the genotyping array. Parameter estimation and imputation were the same as above.

### JPT-1KGP reference panel simulation and imputation

We sampled 7 subsets (size = 100, 500, 1,000, 1,500, 2,000, 2,500, and 3,256) from the JPT_3256_ and then combined the JPT_3256_ with 6 subsets (1KGP-JPT and 1–5 ancestries) of the 1KGP. WGS_993_ was used as the target. The other processing methods were the same as above.

### Number of confident alleles and high-Rsq variants

Haploid dosage (HDS) is the imputed alternative allele’s allelic dosage at the haploid level, which could be obtained by the “--format HDS” option in Minimac4. Briefly, it is in a per-variant and per-individual manner and a higher value represents an imputed alternative allele is more certain. We quantified the θ value’s impact on the imputed dosages by the number of confident alleles (HDS > 0.9). Rsq was used to judge how many high-Rsq (Rsq > 0.7) variants could be passed to the downstream analyses.

Furthermore, we determined how many confident alleles and high-Rsq variants could be obtained only from the additional distant ancestries in the multi-ancestry reference panel. We defined ancestry-specific variants as follows: EUR-only variants only existed in the 1KGP-EUR_403_, and it has no non-EUR variants. JPT_3256_-only and 1KGP-EAS-only variants only existed in the JPT_3256_ and 1KGP-EAS, respectively. Non-EAS variants were not found in the JPT_3256_ or 1KGP-EAS.

### Replication of the TOPMed imputation pipeline

We followed the TOPMed imputation pipeline (Accessed on Oct. 15, 2022, https://topmedimpute.readthedocs.io/) and used the 1KGP and JEWEL3k reference panels. Specifically, the HapMap2 genetic map was used as a reference for the θ value instead of estimating it using Minimac3. WGS_993_ was used as the target. As the θ value was not outputted by default, we modified the Minimac4’s source code to obtain the transformed θ value (**Supplementary Method 5**).

## Results

### BBJ imputation using the four reference panels

The BBJ-180k was imputed using the TOPMed, 1KGP, BBJ1k, and JEWEL3k reference panels (**Figure 1A**). Characteristics of each panel and the target sample are listed in **Table 1**. We categorized the imputed variants by MAF and Rsq in each imputed dataset. With more EAS samples in the panel, more variants with low MAF passed each Rsq threshold (**Figure 2A** and **Supplementary Table 1**). In addition to the absolute number, unique variants were imputed from each panel (**Figure 2B**). These results reproduced the benefits of using large and different reference panels [19].

**Figure 1.**
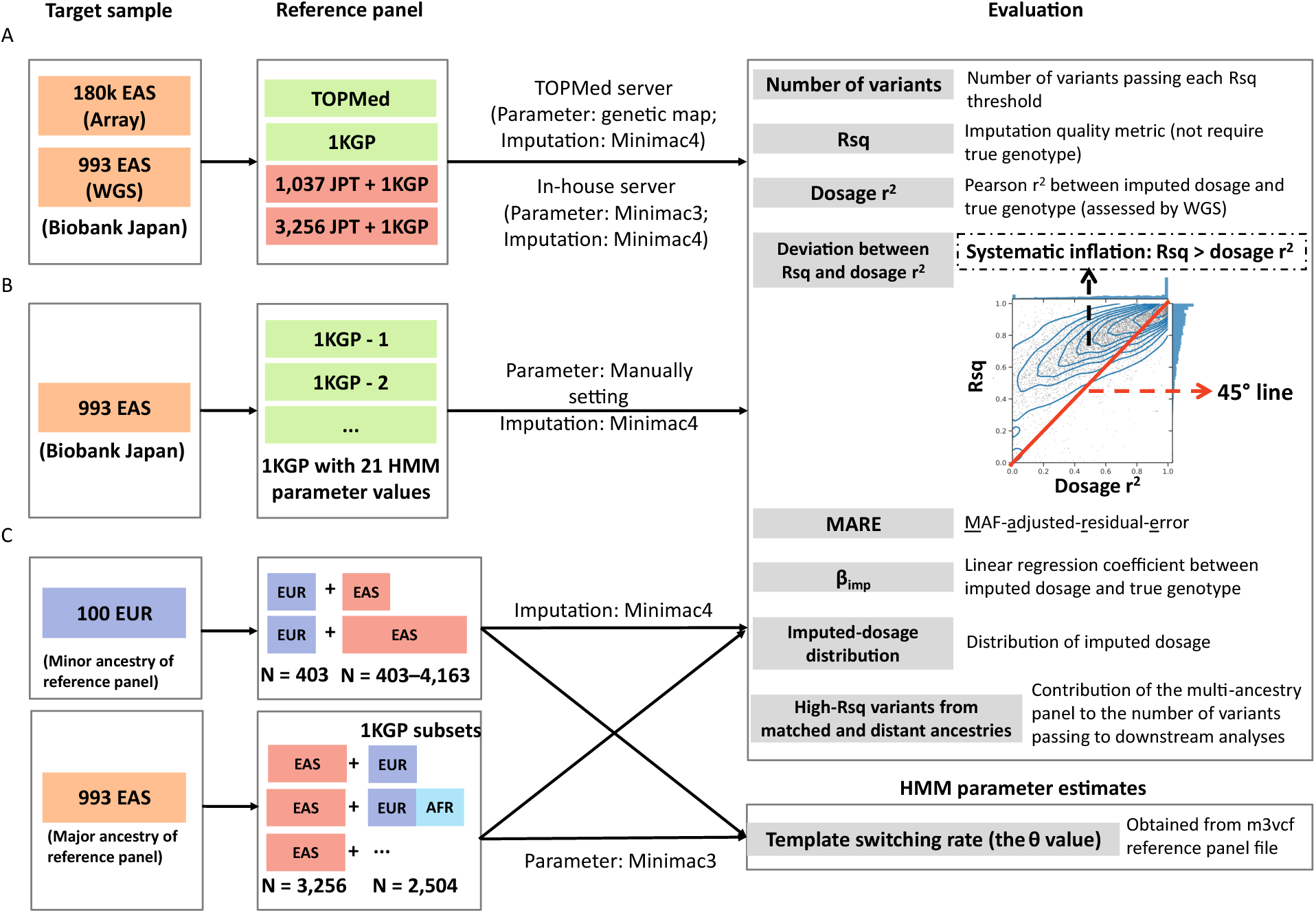
Overview of this study. (A) Imputation using the TOPMed, 1KGP, BBJ1k (1,037 Japanese (JPT) WGS + 1KGP), and JEWEL3k (3,256 JPT WGS + 1KGP) reference panels. The 180k East Asian (EAS) samples in the Biobank Japan (BBJ) first cohort were imputed against the four panels. Samples with whole genome sequencing (WGS_993_) were used to empirically evaluate the imputation performance, including a count of imputed variants and estimated (Rsq) and true (dosage r^2^) imputation accuracy. The deviation between Rsq and dosage r^2^ was examined and quantified analytically. Two novel metrics, MAF-adjusted-residual-error (MARE) and β_imp_, were introduced to indicate the different imputed-dosage distributions underlying the deviation. (B) Simulations to evaluate the relationship between the hidden Markov model (HMM) parameter, Rsq, dosage r^2^, the deviation between Rsq and dosage r^2^, and imputed dosage. The reference panel and target sample were fixed to the 1KGP and WGS_993_, and 21 scaled template switching rates (the θ value) were used for running imputations. Evaluations were the same as in (A). (C) Simulations to evaluate the relationship between the reference panel, HMM parameter, imputation metrics, and imputation result. Two scenarios were simulated to represent the imputations of the major and minor ancestry in a multi-ancestry panel. The 3,256 JPT WGS and 1KGP were used to simulate reference panels with different sizes and ancestral diversities. European (EUR) and JPT samples were used as the target. Imputation results were evaluated the same as in (B). In addition, we evaluated how the Minimac3’s parameter estimates change with the panel composition.

**Table 1.** Characteristics of the reference panels and target samples used in this study, and the number of variants imputed.

| Reference panel | Reference panel size | Size of East Asian subjects | Percentage of EAS subjects | Sequencing depth | Target sample size | Target sample selection criteria | Number of imputed variants on autosomes |
| --- | --- | --- | --- | --- | --- | --- | --- |
| 1KGP | 2,504 | 504 | 20.13% | 4–8 × | 180,882 | All EAS in BBJ -180k passed QC | 47,109,465 |
| BBJ1k | 3,541 | 1,541 | 43.52% | 1KGP + 30 × | 179,947 | Same as above. Exclude samples in the reference panel | 59,387,070 |
| JEWEL3k | 5,760 | 3,760 | 65.28% | 1KGP + 15–30 × | 177,930 | Same as above. Exclude samples in the reference panel | 69,076,783 |
| TOPMed | 97,256 | 1,184 | 1.22% | 38 × | 180,882 | All EAS in BBJ-180k passed QC | 291,214,407 |

**Figure 2.**
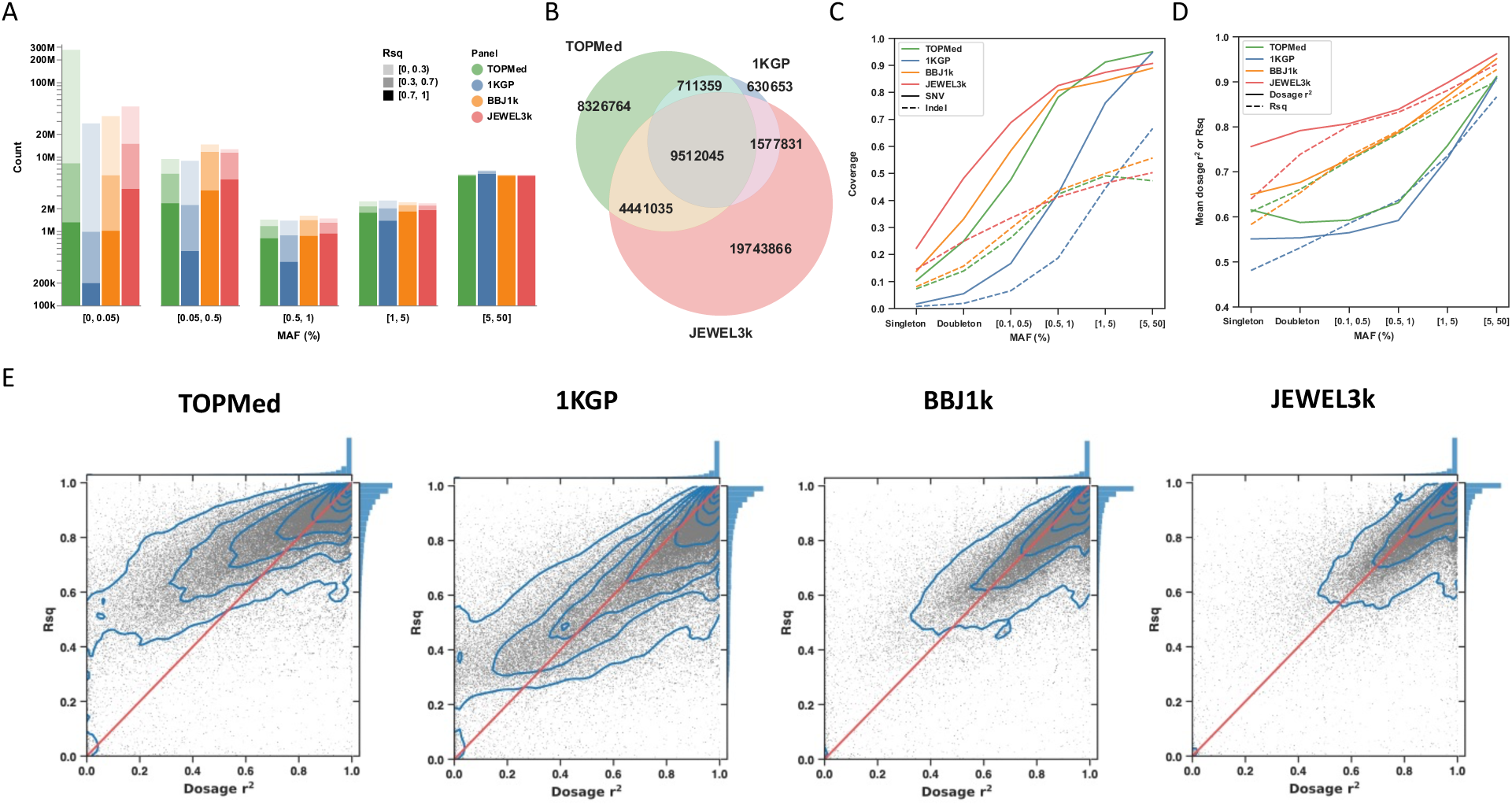
Imputation results using the TOPMed, 1KGP, BBJ1k, and JEWEL3k reference panels. (A) Count of imputed variants using the four panels, stratified by minor allele frequency (MAF) and Rsq in each imputation result. (B) Count of unique and shared variants (with Rsq ≥ 0.3) using the TOPMed, 1KGP, and JEWEL3k panels. (C) Fraction of SNV and indel in WGS_993_ that could be imputed with Rsq ≥ 0.3 (coverage) using each panel, stratified by MAF of WGS_993_. (D) The mean dosage r^2^ and Rsq, stratified by MAF of WGS_993_. (E) Comparison of dosage r^2^ and Rsq in the imputation results using the four panels. The scatter represents the variants, with the blue lines showing the density. The red line shows that Rsq equals dosage r^2^, and the histograms on the side show the distribution. In (C), (D), and (E), variants on chromosome 19 were used. In (D) and (E), overlapping variants imputed by the four panels (Rsq ≥ 0.3 using all panels) were used.

We then used WGS_993_ to empirically evaluate the imputation performance. MAF and MAC of WGS_993_ were used to categorize the variants into six bins: common (MAF ≥ 5%), low-frequency (5% > MAF ≥ 1% and 1% > MAF ≥ 0.5%), rare (0.5% > MAF > 0.1%), doubleton (MAC = 2), and singleton (MAC = 1). For common single nucleotide polymorphisms (SNVs), the TOPMed imputation result showed the highest coverage (0.950), whereas BBJ1k (0.890) and JEWEL3k (0.907) were lower than 1KGP (0.948). This loss of variants was possibly due to the additional QC steps in combining the 1KGP with Japanese WGS (**Supplementary Method 2**). As MAF decreased, the coverage also decreased; however, more EAS samples in the reference panel mitigated this decrease, as expected (**Figure 2C**).

The 155,297 shared SNVs (on chr19) imputed by the four panels were used to compare imputation accuracy. The mean dosage r^2^ using the TOPMed panel was between those of 1KGP and BBJ1k (**Figure 2D**). However, Rsq was substantially upward biased only in the TOPMed imputation result (**Figure 2E**). Using all SNVs and short insertions and deletions (indels) with Rsq ≥ 0.3, although with higher coverage, the TOPMed imputation result showed an inferior dosage r^2^ than 1KGP (**Supplementary Figure 1**).

### Quantifying the deviation between Rsq and dosage r^2^

The deviation between Rsq and dosage r^2^ was persistent in all MAF bins only when using the TOPMed panel (**Supplementary Figure 2**), which indicated a potential systematic bias from the imputation pipeline or the reference panel. To justify our observation, we analytically derived the relationship between Rsq and dosage r^2^ (**Methods**). Two novel metrics, MARE and β_imp_, were introduced to quantify the deviation. MARE, a MAF-adjusted form of residual error, takes a value between 0 and 1, and increases with *SS_res_*. β_imp_ describes the distinguishability between the mean imputed dosage of each true genotype group. Rsq is the ratio between observed and expected variance and dosage r^2^ shows the correlation. Under the assumption of “well-calibration” (the posterior allele probability from imputation equals the expected true allele dose), Rsq equals dosage r^2^ [10]. Our analytical derivation of the relationship between Rsq and dosage r^2^ did not assume the “well-calibration” (**Supplementary Figure 3–7** and **Supplementary Note 1**). Any two of Rsq, dosage r^2^, MARE, and β_imp_ could entirely quantify the relationship between the imputed dosage and true genotype and determine the other two metrics (Equations 6–7), whereas Rsq or dosage r^2^ alone could not.

To exhibit the relationships, we plotted the theoretical values of MARE and β_imp_ on the coordinates of Rsq and dosage r^2^ (**Figure 3A** and **3F**). It clearly showed that the overestimated Rsq was accompanied by a higher MARE (**Figure 3A**). We used rs142572000 as an example (**Figure 3B–E**). In the TOPMed imputation result, imputed genotypes were more certain (defined as the imputed dosage closer to 0, 1, or 2) (**Figure 3B**) [37], compared to the other three panels (**Figure 3C–E**). The high certainty increased Var(***y***) and Rsq. In **Supplementary Note 4**, the positive relationship between imputed-genotype certainty and Rsq is demonstrated. As discussed further below, high certainty or Rsq did not mean the imputation is more accurate. As shown in **Figure 3B**, many heterozygotes were incorrectly imputed with a dosage of approximately 0 in the TOPMed imputation, causing higher MARE and Rsq, and an even low dosage r^2^.

**Figure 3.**
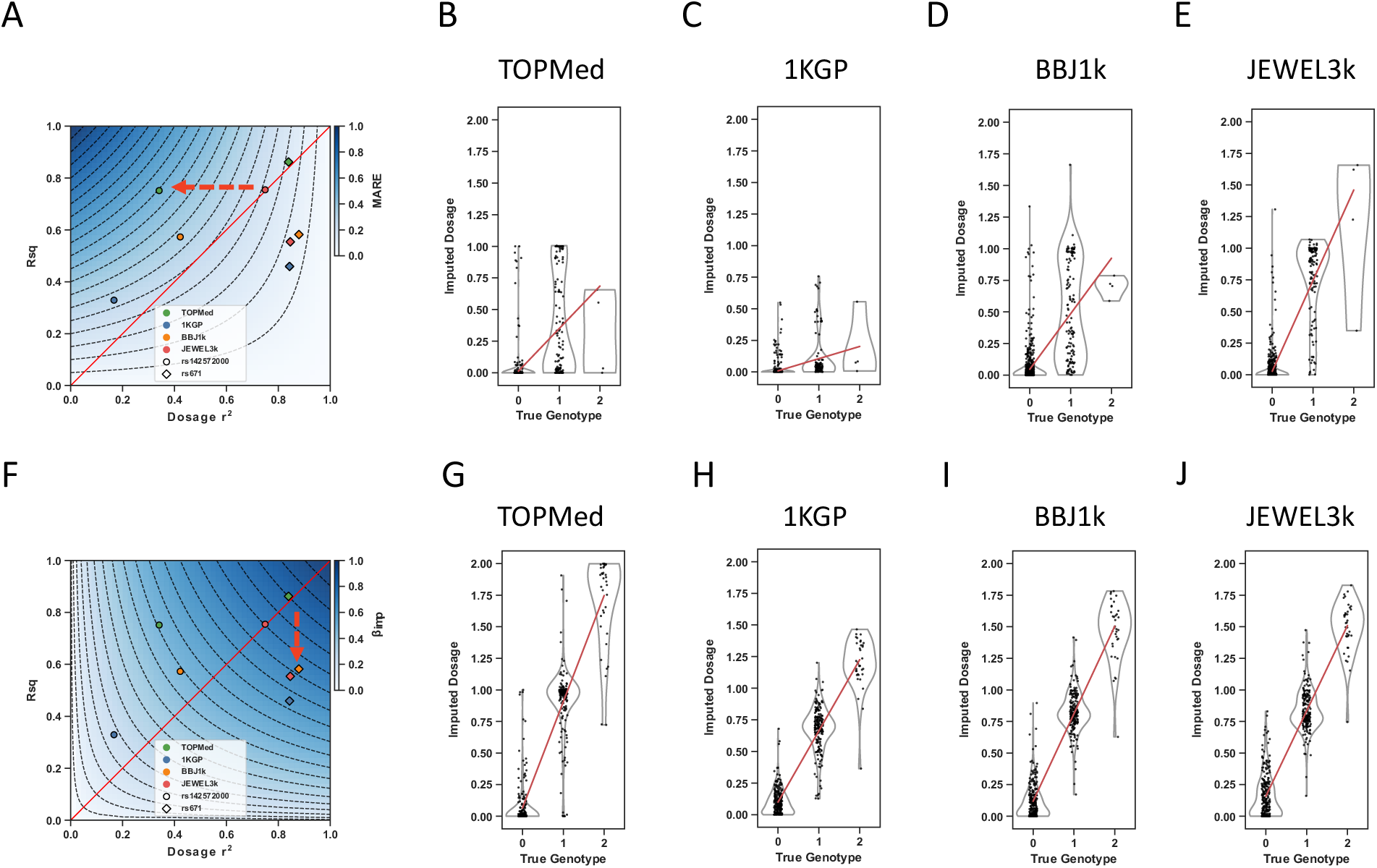
MARE, β_imp_, Rsq, dosage r^2^, and imputed dosage of rs142572000 and rs671. (A) The plot shows Rsq and dosage r^2^ of rs142572000 and rs671, and the theoretical value of MARE on the Rsq ∼ dosage r^2^ plot. (B–E) The plot shows the imputed dosage of rs142572000 in the imputation result using the (B) TOPMed, (C) 1KGP, (D) BBJ1k, and (E) JEWEL3k reference panels. (F) The plot shows the theoretical value of β_imp_ on the Rsq ∼ dosage r^2^ plot. (G–J) The plot shows the imputed dosage of rs671 in the imputation result using the (G) TOPMed, (H) 1KGP, (I) BBJ1k, and (J) JEWEL3k panels. In (A) and (F), the red line shows that Rsq equals dosage r^2^, and the color keys show the theoretical MARE or β_imp_ values. The dashed lines represent the contours of MARE or β_imp_, and adjacent contours have a value difference of 0.05. The red dashed arrow highlights the difference in MARE and β_imp_ (crossing how many contour lines) for variants showing a similar Rsq but different dosage r^2^ (rs142572000), or similar dosage r^2^ but different Rsq (rs671). In (B–E) and (G–J), the strip plot shows the individual and the violin plot shows the distribution. The red line shows the linear regression line between the imputed dosage and true genotype. In (B– E), MARE is 0.473, 0.274, 0.328, and 0.187, and β_imp_ is 0.336, 0.097, 0.439, and 0.717. In (G–J), MARE is 0.139, 0.072, 0.071, and 0.085, and β_imp_ is 0.843, 0.565, 0.694, and 0.675.

Variants with dosage r^2^ < Rsq showed a lower β_imp_ (**Figure 3F**). We used rs671 as another example (**Figure 3G–J**). Except for the TOPMed imputation result, imputed dosages were shrunk to the AAF in EAS (0.247) (**Figure 3H–J**), causing uncertainty in the imputed genotype, and decreased β_imp_, MARE, and Rsq. However, imputed dosages were still highly correlated with the true genotypes (**Figure 3F**), causing dosage r^2^ > Rsq. Taking the two examples together, imputation results of a similar Rsq or dosage r^2^ may have a great difference in the imputed dosage, thus, more comprehensive evaluations are necessary and the deviation between Rsq and dosage r^2^ should not be ignored.

We categorized MARE into Rsq bins and β_imp_ into dosage r^2^ bins to compare them between reference panels and to the expected values when assuming Rsq equals dosage r^2^. The mean MARE of the TOPMed result was above the expected for Rsq bins 0.35–0.9, and the mean β_imp_ of the 1KGP result was below those of the other panels (**Supplementary Figure 8**). These results suggested that the TOPMed result was more certain and might contain more wrongly imputed genotypes (**Figure 3B** and **3G**), while the 1KGP result might possess higher shrinkage in the imputed dosage, as shown above (**Figure 3C** and **3H**).

### Template switching rate impacts the deviation between Rsq and dosage r^2^

The high certainty suggested an overconfident matching between the reference panel and target sample. We investigated how the template switching rate (θ) used in the HMM affects the imputed-allele certainty using the 1KGP reference panel and 21 scalings of the θ value (0.01–100-fold of that estimated by Minimac3; **Methods**; **Figure 1B**). Low θ values caused Rsq > EmpRsq (EmpRsq is the alternate of dosage r^2^; **Methods**) (**Supplementary Figure 9A–C**), while high θ values caused Rsq < EmpRsq (**Supplementary Figure 9E–G**). Using rs10410162 as an example, Rsq and MARE increased as the θ value decreased (**Figure 4A**), with increasing certainty of the imputed alleles (**Supplementary Figure 10A–C**) and the deviation toward Rsq > EmpRsq (**Figure 4B**). As the θ value increased, the imputed allelic dosages were shrunk to the AAF (0.351) (**Supplementary Figure 10E–G)**, resulting in a more drastic decrease in Rsq than EmpRsq (**Figure 4A**), thereby causing EmpRsq > Rsq. Notably, EmpRsq and β_imp_ were roughly maintained unless the θ value was scaled up by 2-fold or higher (**Figure 4A** and **4C**), suggesting EmpRsq was insensitive to altering the θ value, particularly the downscaling. In contrast, Rsq was sensitive to the θ value and consistently increased with downscaling of the θ value, leading to the deviation between Rsq and EmpRsq.

**Figure 4.**
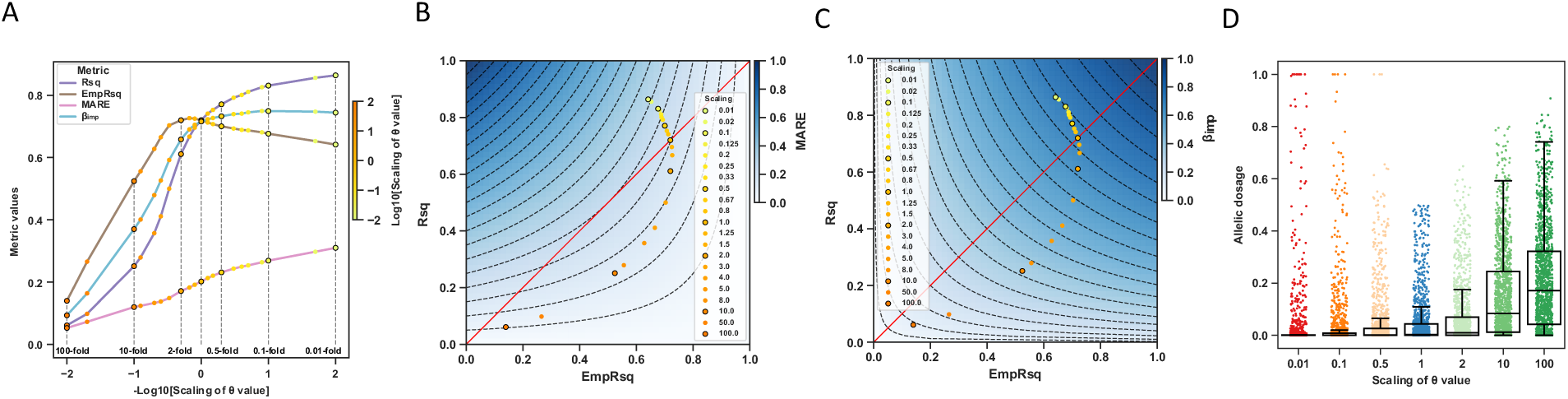
Changes in the metrics value and the imputed allelic dosage with different scalings of the θ value. (A) Rs10410162 is used as an example to visualize the changes in Rsq, EmpRsq, MARE, and β_imp_ (y-axis) with the 21 scalings of the θ value (x-axis). To have a better illustration, the x-axis is in the minus log scale. The scalings of 100, 10, 2, 1, 0.5, 0.1, and 0.01-fold are circled, marked with vertical dashed lines, and labeled. The other scaling folds are shown as the scatters, with the color key denoting the fold in a log scale. (B–C) Rs10410162 is used as an example to visualize the deviation between Rsq, EmpRsq, and MARE (B) or β_imp_ (C). The red line shows that Rsq equals dosage r^2^, and the color keys show the theoretical MARE or β_imp_ values. The dashed lines represent the contours of MARE or β_imp_, and adjacent contours have a value difference of 0.05. The 21 scatters are the same as (A). (D) The plot shows the imputed allelic dosage of a randomly selected target haplotype (chr19) with the 7 scalings of the θ value. The scatters show variants. The boxes show the median, upper (75%), and lower (25%) quartiles. The whiskers show the 1.5-fold of the interquartile range (IQR) extended from the upper or lower quartile if a value exceeds them; otherwise, they show the maximum or minimum value. Only variants with imputed allelic dosage < 0.5 and 0.3 < EmpRsq < 0.8 when using the 1-fold θ value are shown.

We also showed that low θ values increased the imputed-allele certainty using a randomly selected target haplotype **(Figure 4D** and **Supplementary Figure 11).** High certainty might cause some variants to be wrongly imputed as the opposite allele (**Supplementary Figure 12**). An explanation is provided in **Supplementary Note 5**.

Such imputed-dosage properties were also revealed by the mean MARE and β_imp_ across all variants on chr19 (**Supplementary Figure 13**). Taken together, the simulated imputation using an extremely low θ value mimics the high certainty and Rsq overestimation in the TOPMed imputation result in EAS (**Figure 3**).

### Template switching rate impacts the imputation performance

Using the modified 1KGP panels, although the mean EmpRsq was insensitive to the scaling of the θ value (maximum difference of 0.034 for the θ value scaling between 0.01 and 2-fold; **Figure 5**), low θ value increased Rsq (**Figure 5**). We evaluated the number of confident alleles (HDS > 0.9) and high-Rsq variants (Rsq > 0.7) (**Methods**). Downscaling the θ value increased the number of confident alleles and high-Rsq variants (**Supplementary Table 2**). When the θ value was 0.5-fold, the number of confident alleles and high-Rsq variants increased by 5.46% and 8.89%, respectively, and if the θ value was 2-fold, these numbers decreased by 11.8% and 13.8%, respectively. These results indicated that the θ value shaped the imputed allelic dosage and changed Rsq and the number of high-Rsq variants while leaving EmpRsq almost the same.

**Figure 5.**
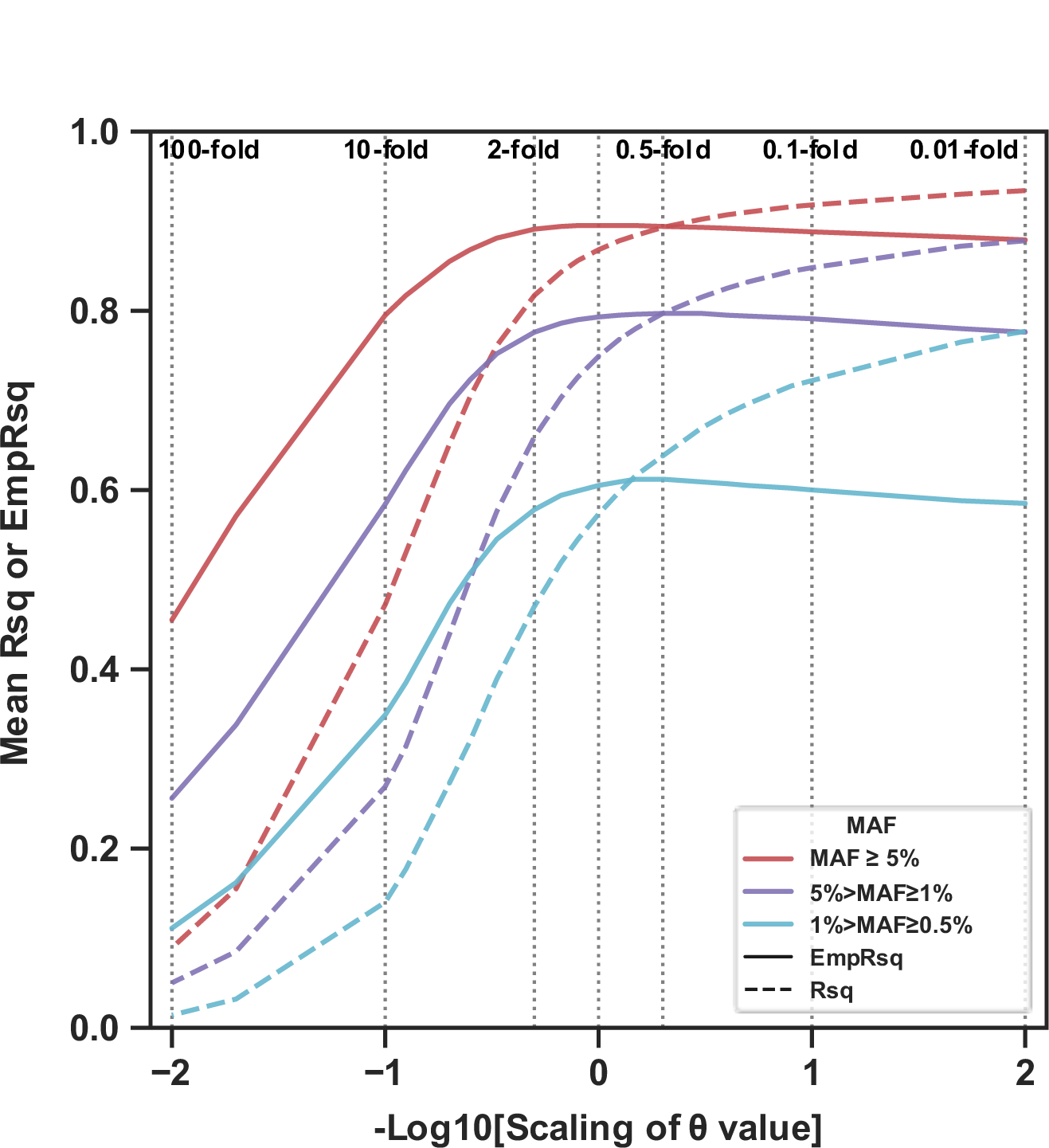
Rsq and EmpRsq using the 1KGP panel with different scalings of the θ value. The plot shows the mean Rsq and EmpRsq of each imputation result, stratified by the MAF of the array. The scalings of 100, 10, 2, 1, 0.5, 0.1, and 0.01-fold are marked with vertical dashed lines and labeled.

### Reference panel and θ estimates

We evaluated how Minimac3’s parameter estimation changed with the composition of the reference panel (**Supplementary Method 4**, **Supplementary Figure 14**, and **Supplementary Note 6**). The θ estimates decreased with the sample size of a single ancestry and increased with ancestral diversity when size was fixed (**Figure 6A** and **6B**). On pooling samples from different ancestries together at a small panel size, the θ estimates still decreased as the panel size increased (**Figure 6C**); however, at a larger panel size, it increased with simultaneously increasing the panel size and ancestral diversity (**Figure 6D**), suggesting that the θ estimates were in a trade-off between the panel size and ancestral diversity (**Figure 6E**). **Supplementary Note 6** explains these effects.

**Figure 6.**
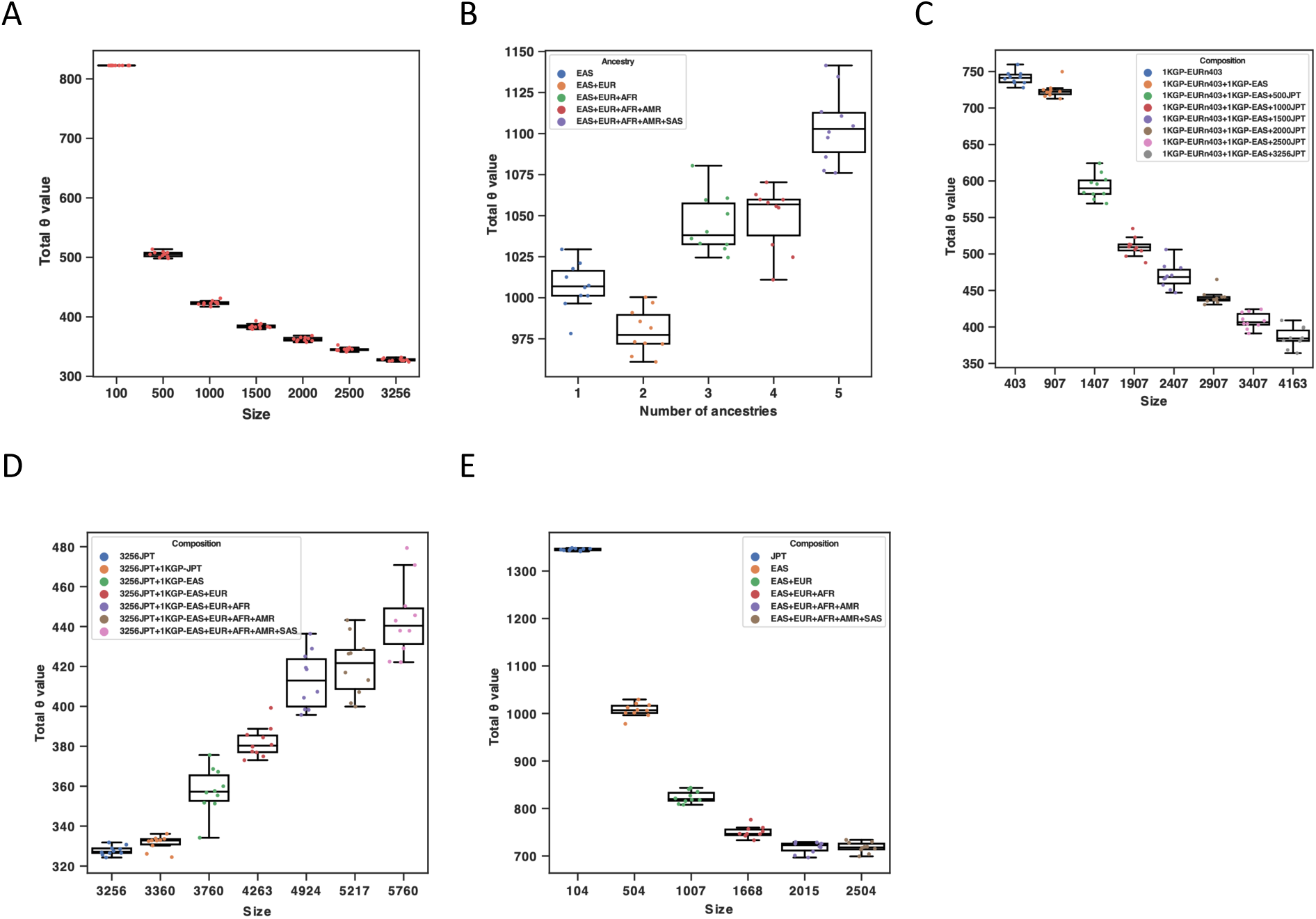
Total θ value of the simulated reference panels. The Supplementary Figure hows the total θ value of simulated reference panels, estimated by Minimac3. (A) JPT-only panels with different sample sizes. (B) Fixed-size panels (504) with 1–5 levels of ancestral diversity, as shown in the legend. (C) Adding large-size (504–3,760) EAS samples to a small-size (403) EUR panel. (D) Adding 1KGP samples to a large-size (3,256) JPT panel. (E) Simultaneously increasing the size and ancestral diversity using 1–5 ancestries in the 1KGP. In each plot, the scatters show the values of 10 runs. The boxes show the median, upper (75%), and lower (25%) quartiles. The whiskers show the 1.5-fold interquartile range (IQR) extended from the upper or lower quartile if a value exceeds them; otherwise, they show the maximum or minimum value. In (C), EURn403 represents the 403 EUR. In (C) and (D), 500–3256JPT represents the number of JPT samples in the panel.

### Fitness of the θ value, deviation, and imputation performance

To elucidate the multi-ancestry reference panel’s impact on the imputation result, we simulated two scenarios using the multi-ancestry reference panels: the target sample was from the (1) minor and (2) major ancestry (**Figure 1C**).

#### Scenario 1: The target sample was from the minor ancestry

We simulated 8 EUR-EAS reference panels and used 100 EUR samples as the target (**Methods**). As the panel size increased and θ value decreased (**Table 2**), Rsq and MARE were upwardly biased, as expected (**Supplementary Figure 15** and **S16**). The mean EmpRsq decreased consistently with the addition of EAS samples to the reference panel (**Table 2**). However, only a marginal difference was observed (maximum difference of 0.020).

**Table 2.** Reference panel description, the total θ value estimated by Minimac3, the mean EmpRsq, and the mean Rsq using the simulated EUR-EAS reference panels with different sizes and EUR proportions, and 100 EUR as the target sample.

| Reference panel | Reference panel size | EUR (%) | Total $\theta$ value | MAF ≥ 5% | | 5% ≥ MAF > 1% | |
| --- | --- | --- | --- | --- | --- | --- | --- |
|  |  |  |  | EmpRs<sub>q</sub> | Rs<sub>q</sub> | EmpRs<sub>q</sub> | Rs<sub>q</sub> |
| 1KGP-EURn403 | 403 | 100 | 740.0 | 0.849 | 0.797 | 0.75 | 0.679 |
| 1KGP-EURn403+1KGP-EAS | 907 | 44.43 | 722.7 | 0.841 | 0.795 | 0.743 | 0.7 |
| 1KGP-EURn403+1KGP-EAS+500JPT | 1,407 | 28.64 | 595.9 | 0.839 | 0.814 | 0.744 | 0.736 |
| 1KGP-EURn403+1KGP-EAS+1000JPT | 1,907 | 21.13 | 534.9 | 0.836 | 0.822 | 0.74 | 0.752 |
| 1KGP-EURn403+1KGP-EAS+1500JPT | 2,407 | 16.74 | 466.0 | 0.834 | 0.831 | 0.739 | 0.767 |
| 1KGP-EURn403+1KGP-EAS+2000JPT | 2,907 | 13.86 | 437.5 | 0.833 | 0.835 | 0.737 | 0.774 |
| 1KGP-EURn403+1KGP-EAS+2500JPT | 3,407 | 11.83 | 408.4 | 0.831 | 0.839 | 0.734 | 0.78 |
| 1KGP-EURn403+1KGP-EAS+3256JPT | 4,163 | 9.68 | 364.1 | 0.829 | 0.844 | 0.732 | 0.788 |

EURn403 represents the 403 EUR; 1KGP-EAS represents the 504 EAS; 500–3256JPT represents the number of JPT samples in the reference panel. Minor allele frequency (MAF) is determined by the genotyping array.

There were 114,606 EUR-only and 696,017 non-EUR variants (**Methods**). As the panel size increased and θ value decreased, the number of confident alleles increased from 84,797 to 101,116 and from 0 to 458 for EUR-only and non-EUR variants, respectively (**Supplementary Table 3**). The number of high-Rsq variants increased from 163,468 to 200,161, 18,646 to 27,941, and 0 to 521 for all, EUR-only, and non-EUR variants, respectively (**Supplementary Table 3**). These results revealed that a lower θ value increases the number of confident alleles and high-Rsq variants, without improving the EmpRsq (**Supplementary Table 3**). Meanwhile, even if the reference panel comprised 90.3% EAS samples, non-EUR variants only comprised 0.26% of the total variants in the imputation result (when setting a cutoff of Rsq > 0.7). The majority of variants gaining from the larger panel when imputing the under-represented ancestry was because of the lower θ value.

#### Scenario 2: The target sample was from the major ancestry

We simulated 13 JPT-1KGP reference panels and used WGS_993_ as the target (**Methods**). When only JPT samples were in the panel, the mean EmpRsq and Rsq increased with the panel size, as expected (**Supplementary Figure 17**). When combined with the 1KGP subsets, the mean EmpRsq was highest when using JPT_3256_+1KGP-EAS for variants with MAF ≥ 1% and JPT_3256_+1KGP-JPT for variants with 1% > MAF ≥ 0.5% (**Table 3**). Adding other ancestries decreased the mean EmpRsq marginally (maximum difference < 0.01 for all MAF categories). The mean Rsq was the highest when using JPT_3256_ and decreased with the addition of more ancestries, with maximum differences of 0.007, 0.024, and 0.048 for variants with MAF ≥ 5%, 5% > MAF ≥ 1%, and 1% > MAF ≥ 0.5%, respectively (**Table 3**). None of the imputation results showed a noticeable deviation in the MARE and β_imp_. However, the MARE and β_imp_ of the JPT_3256_, JPT_3256_+1KGP-JPT, and JPT_3256_+1KGP-EAS results were closer to the expected values (**Supplementary Figure 18**), while adding other ancestries made MARE and β_imp_ correspond to the condition of using a higher θ value.

**Table 3.**
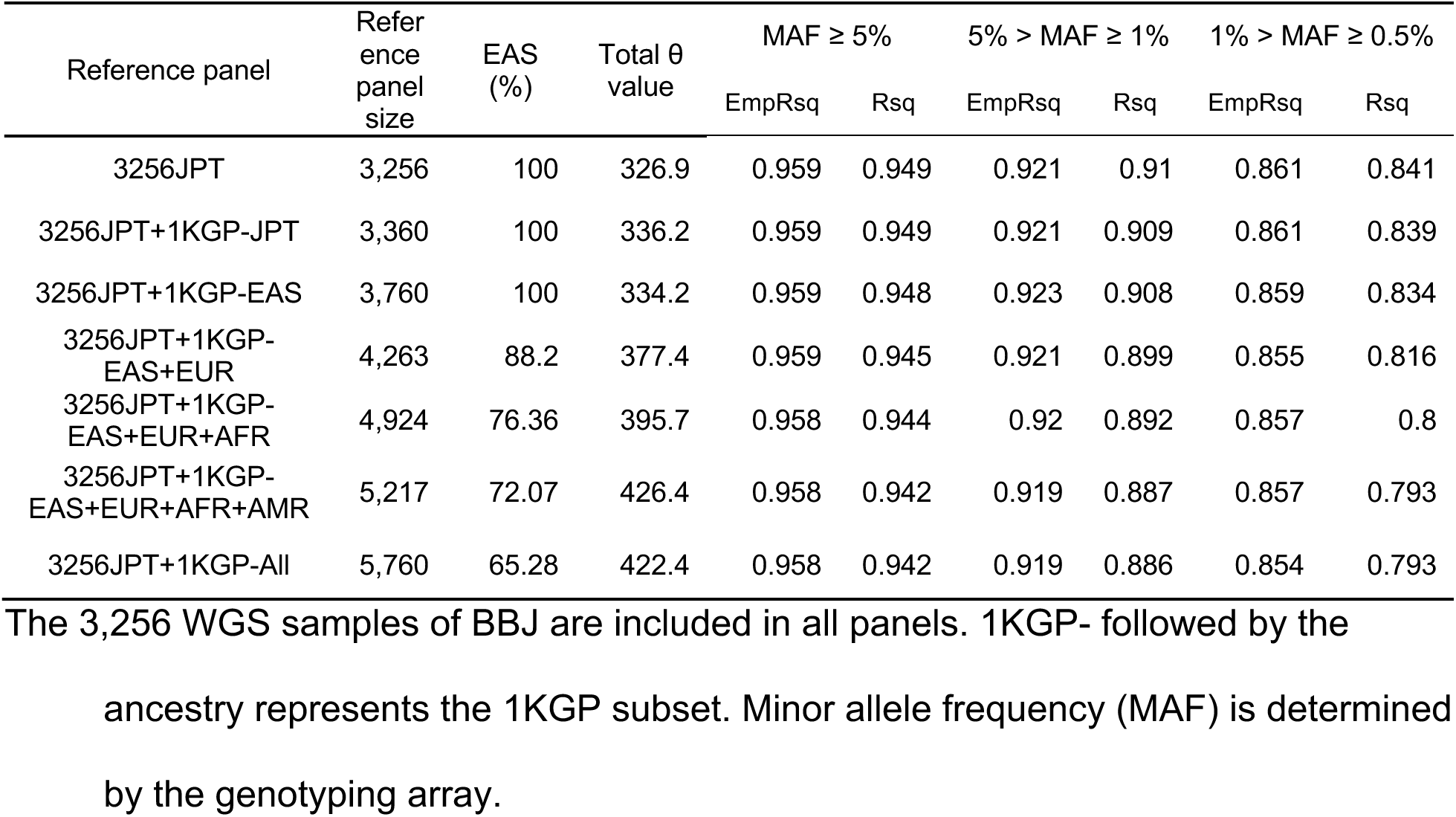
Reference panel description, the total θ value estimated by Minimac3, the mean EmpRsq, and the mean Rsq using the simulated JPT-1KGP reference panels with different sizes and ancestral diversity, and WGS_993_ as the target sample.

There were 74,490 JPT_3256_-only, 7,627 1KGP-EAS-only, and 495,489 non-EAS variants (**Methods**). From JPT_3256_ to JPT_3256_+1KGP, the number of confident alleles decreased from 24,599 to 23,666, increased from 0 to 131, and increased from 0 to 254 for these three groups of variants (**Supplementary Table 4**). The number of high-Rsq variants decreased from 274,343 to 264,221 (a decrease of 3.69%) when using JPT_3256_ or JPT_3256_+1KGP (**Supplementary Table 4**). Only 265 non-EAS variants reached an Rsq > 0.7 when using JPT_3256_+1KGP, which was 0.10% of the total variants. Hence, combining with the 1KGP caused fewer confident alleles and high-Rsq variants to remain in the imputation results, as expected from using higher θ values. Both scenarios suggest that the distant ancestry in the reference panel affects the θ estimates and the number of high-Rsq variants. However, the expanded haplotypes and variant sets from distant ancestries provided only a few additional variants.

### EAS imputation using the public reference panels

We used the TOPMed imputation pipeline (**Methods**) and observed an upward deviation of Rsq, particularly when using the 1KGP panel (**Supplementary Figure 19**). The θ value transformed from the genetic map was 0.15- and 0.27-fold of that estimated by Minimac3 when using the 1KGP and JEWEL3k panels, respectively (**Supplementary Table 5**).

As the θ estimates decreased with the size of the major ancestry (**Figure 6C**), it would be underestimated for the minor ancestry in the multi-ancestry panel regardless of using the genetic map or estimating it by the Mimimac3. One example was the HRC panel, which did not use the genetic map but induced Rsq overestimation in the BBJ-180k (**Supplementary Figure 20**). Thus, caution may be required when imputing the under-represented population using large ancestry-imbalanced panels under the current framework.

## Discussion

The deviation between Rsq and dosage r^2^ in the imputation result is raised from the imputed dosage, and has been widely observed using different reference panels and software [9,15,38]. One reason was that the θ value used in the HMM does not fit the panel size, ancestral components in the reference panel, and the target population. When using the multi-ancestry reference panel, distant ancestries affect the θ estimates (in Minimac3/4), then the imputed dosage and Rsq. Our simulations indicated that the lower θ value used by the larger panel could increase the imputed-genotype certainty and the number of high-Rsq variants but not dosage r^2^. It may cause confusing benchmarking results and increase the chance of false positives in association tests. Ferwerda et al. reported height and body mass index association signals in an ethnically diverse cohort only when imputing against the TOPMed panel [39]. Bai et al. reported the highest Rsq but lowest dosage r^2^ when imputing the Han Chinese using the HRC panel, compared to the 1KGP and a population-specific panel [22], similar to what we had observed in Japanese. In addition to simply comparing dosage r^2^ or Rsq, we provided the changes in those metrics, the θ estimates, imputed-dosage certainty, and the number of high-Rsq variants passing to downstream analyses. We provided the script (https://github.com/shimaomao26/impumetric) to allow users to check their results using the leave-one-out imputation of Minimac4 (external WGS not required) or additional WGS data.

The TOPMed imputation pipeline used the θ value transformed from the HapMap2 genetic map by assuming 0.01 centimorgan (cM) corresponds 1% switching rate. The switching rate would thereby be fixed given the variant’s base pair position and cM. Two reasons may explain why it works well. First, the genetic map did not significantly change the imputation accuracy, as verified in a Finnish study [40]. We had also shown that dosage r^2^ was insensitive to the θ value. Second, the θ value was only underestimated for the EAS (size = 1,184), but the sizes of EUR, AFR, and AMR (size = 17,085–47,159) might fit this value. Previous studies reported that Rsq in the TOPMed imputation result was sometimes misleadingly high in AFR [38] and even EUR [41]. A recent paper reported that the TOPMed panel was more robust to the low-density genotyping array than the HRC and 1KGP panels [18]. These results also suggested that a fixed low θ value might be used by the TOPMed pipeline, as we have inferred. Further studies are warranted to comprehensively elucidate the impacts on genetic studies.

Our results suggest that avoiding ancestral diversity is best when more than 3,000 WGS samples are available to construct a JPT reference panel. Increased diversity would then only have a marginal impact on dosage r^2^ but would affect the individual’s imputed dosage and cause fewer variants to pass a predetermined Rsq filter. Zhang et al. and Cong et al. similarly observed that adding the 1KGP to about 3,000 Chinese WGS samples would neither benefit nor harm dosage r^2^ [26,27]. We focused on explaining the reason in this work. Further studies are warranted to take advantage of combining population-specific and public WGS datasets while avoiding our identified problems.

Our study has several limitations. First, our simulations of the reference panel were study-specific. As discussed, the θ estimates decreased with the panel size and increased with the ancestral diversity. However, a larger panel typically increases the diversity simultaneously. Therefore, the θ value and imputation result need to be discussed case by case. This limitation also implies that the general experience may not work well for a new reference panel and target dataset. Second, in the simulations of multi-ancestry reference panels (scenarios 1 and 2), WGS datasets were merged using IMPUTE2 [9]. We did not investigate the impact of IMPUTE2 but treated the merged dataset similarly to the WGS dataset. It might underestimate the number of high-Rsq variants from the distant ancestry (**Supplementary Note 7)**. Our results using the 1KGP (not modified by IMPUTE2) also revealed that only a limited number of confident alleles and high-Rsq variants in the imputation result were from the distant ancestry (**Supplementary Note 7** and **Supplementary Table 2**). Therefore, the conclusion that distant ancestry in the reference panel affects the θ estimates rather than providing high-Rsq variants would be valid. However, further studies should be conducted to determine the impact of the panel-merging method and the net gain of variants from distant ancestries.

In summary, we explained that the HMM parameter could be a potential reason for inaccurate Rsq and inferior dosage r^2^ when using large multi-ancestry reference panels. This is also the first study of the relationship between the template switching rate, imputed-genotype certainty, Rsq, and dosage r^2^. We envision that our methods and conclusions could provide insights into benchmarking studies, construction of reference panels, and development of imputation algorithms and pipelines in the future.

### Funds

This research was supported by the Ministry of Education, Culture, Sports, Sciences and Technology (MEXT) of Japanese government and the Japan Agency for Medical Research and Development (AMED) under grant numbers JP18km0605001 (the BioBank Japan project) and JP19km0405215 (to C.T., K.M., and Y.K.).

## Acknowledgments

We thank the staff of the BBJ for collecting and managing samples and clinical information. We acknowledged the Human Genome Center, the Institute of Medical Science, the University of Tokyo (http://sc.hgc.jp/shirokane.html) for providing the super-computing resources.

## Web resources

Impumetric, https://github.com/shimaomao26/impumetric

TOPMed imputation server, https://imputation.biodatacatalyst.nhlbi.nih.gov/#!

Michigan Imputation server, https://imputationserver.sph.umich.edu/index.html

Minimac3, https://genome.sph.umich.edu/wiki/Minimac3

Minimac4, https://genome.sph.umich.edu/wiki/Minimac4

EAGLE, https://alkesgroup.broadinstitute.org/Eagle/

Bcftools, https://samtools.github.io/bcftools/

LiftOver, https://genome.ucsc.edu/cgi-bin/hgLiftOver

## Data and code availability

We provided the tool to replicate the metrics used in this work (https://github.com/shimaomao26/impumetric). Other scripts were also deposited there. Genotype data for BBJ and the TOPMed imputation results were deposited at NBDC Human Database (research ID: hum0014 and hum0311, respectively).

## Appendix

### Imputation quality metrics

Rsq: The standard quality metric in Minimac. The ratio between observed variance and expected variance of imputed allelic dosage.

INFO: The standard quality metric in IMPUTE. The ratio between observed information and complete information of imputed genotype distribution.

Dosage r^2^: The squared Pearson correlation between imputed dosage and true genotype.

EmpRsq: The squared Pearson correlation between imputed allelic dosage and true allele dose. Provided by Minimac and is only available for the genotyped markers on array.

MARE: The minor allele frequency adjusted residual error in linear regression between imputed dosage and true genotype.

β_imp_: The regression coefficient between imputed dosage and true genotype.

## Notes

### Competing Interest Statement

The authors have declared no competing interest.

### Summary of Updates

Title, abstract, and discussion were updated to clarify this study's focus on imputation accuracy and metric value when using multi-ancestry reference panels.

