## Supplemental Material for "Genotype imputation accuracy and the quality metrics of the minor ancestry in multi-ancestry reference panels"

### Supplementary Methods

#### Supplementary Method 1. Cohort specification of the Biobank Japan (BBJ), genotyping, QC, and selection of EAS subjects

The BBJ project first cohort enrolled 200k participants from 2003 to 2007 [1,2]. Subjects were genotyped by either the Illumina HumanOmniExpressExome BeadChip or a combination of the Illumina HumanOmniExpress and HumanExome BeadChips. The coordinates were on the genome build hg19. QC steps were described in the previous literature [3]. Briefly, variants with (1) call rate < 99%; (2) p-value for Hardy Weinberg equilibrium (HWE) < 1e-6; (3) number of heterozygotes < 5 were excluded. Additional QC was performed by comparing the genotypes between the whole genome sequencing (WGS) and the array of 939 subjects, and then variants with a concordance rate < 99.5% or a non-reference discordance rate ≥ 0.5% were excluded, as described previously. After QC, 520,378 variants on autosomes remained. Samples with a call rate < 98% were excluded. The ancestries were determined by projecting the principal component (PC) onto the PC space of the 1KGP, and 180,882 individuals of the EAS ancestry were used in this study. We named it BBJ-180k.

#### Supplementary Method 2. Methods to construct the BBJ1k and JEWEL3k reference panels

Methods to construct the BBJ1k reference panel were described in the previous literature [3]. Briefly, 1,037 samples from the BBJ were sequenced at 30 × depth. After calling the variants using GATK (v3.2-2) and performing QC, variants located at the low-complexity regions (LCR; accessed from ftp://ftp.1000genomes.ebi.ac.uk/vol1/ftp/release/20130502/supporting/low_complexity_regions/hs37d5-LCRs.20140224.bed.gz) were removed. BEAGLE was used to impute the missing genotypes. To combine the WGS dataset with the 1KGP, variants located at the multi-allelic sites were removed. Then SHAPEIT was used for phasing, and IMPUTE2 was used to combine them. Singletons and variants at multi-allelic sites were removed from the final merged reference panel. Methods to construct the JEWEL3k reference panel were described in the previous literature [4]. Briefly, a total of 1,491 and 1,765 samples from BBJ were sequenced at 30 × and 15 × depth, respectively. The same process with the BBJ1k was used to construct the JEWEL3k reference panel. One sample in BBJ1k was excluded from the JEWEL3k due to the joint QC.

**Supplementary Method 3. QC, liftover, and variant comparison using the external WGS dataset**

The 1,007 samples with WGS were used to evaluate the imputation quality. Peripheral blood-derived genomic DNA was extracted, prepared by TruSeq Nano DNA Library Preparation Kit, and sequenced at 15 × depth using Illumina HiSeq 2500 (150bp paired-ends). Genotypes were called using BWA-MEM (v0.7.13) and GATK (v3.8). The reference genome was hg19 (build37+decoy from the 1KGP; ftp://ftp-trace.ncbi.nih.gov/1000genomes/ftp/technical/reference/phase2_reference_assembly_sequence/hs37d5.fa.gz). We applied the quality filter of GQ ≥ 20 and DP ≥ 2. Variants with a missing rate > 10% were removed. To avoid the bias caused by the cryptic relatives between the reference panel and target sample, PLINK v1.9 was used to calculate the PI_HAT (using variants on the array which were pruned by Plink with "--indep-pairwise 200 50 0.35" and with MAF ≥ 10%) in a combination of the JEWEL3k and 1,007 WGS samples. No pairs were above 0.1. Fourteen samples were excluded as not in all imputed datasets. We named it $WGS_{993}$. Chromosome 19 was selected as the representative as no variants were on the known inverted regions (by comparing to the UCSC LiftOver chain file; https://hgdownload.soe.ucsc.edu/goldenPath/hg38/liftOver/) between hg38 and hg19. Only variants that could be lifted interchangeably (not to the alternative contig or duplicated position) between hg19 and hg38 were retained. There were 581,172 SNVs and 65,157 indels remaining. For SNVs, we compared the alternative allele frequency (AAF) between the TOPMed imputation result and $WGS_{993}$, then swapped the ref/alt alleles if the AF discordance was less than 15% after swapping (except the palindromic variants). For multi-allelic SNVs and all indels, only variants with the exact ref/alt allele matching and AAF discordance less than 15% were retained. Multi-allelic indels were removed from all imputation results before the comparison.

**Supplementary Method 4. Reference panel simulation and the θ value estimation**

It was unclear how the reference panel affects the θ estimates. To comprehensively investigate the impact of the reference panel on the θ value, we simulated the following five scenarios.

Scenario 1 (Size of a JPT population): We took 100, 500, 1,000, 1,500, 2,000, 2,500, and 3,256 samples from the $JPT_{3256}$. The larger subset always contained all the samples in the smaller subset.

Scenario 2 (Ancestral diversity in a fix-size panel): We extracted 1–5 ancestries from the 1KGP. Then, we shuffled and downsampled the subsets to a size of 504 ten times. We did not maintain consistent ancestry ratios in the downsampled files.

Scenario 3 (Adding large-size EAS samples to a small-size EUR panel): We randomly sampled 403 individuals from the 503 1KGP-EUR samples, combined with the 1KGP-EAS (size = 504) and the six subsets (size = 500–3,256) of the $JPT_{3256}$ made in Scenario 1.

Scenario 4 (Adding 1KGP samples of distant ancestries to a large-size EAS panel): We combined the $JPT_{3256}$ with the 1KGP-JPT and 1–5 ancestries in the 1KGP.

Scenario 5 (Simultaneously increase the panel size and ancestral diversity): We extracted 1–5 ancestries from the 1KGP.

Then, we shuffled the samples in each dataset 10 times. Since the dataset had already been cleaned and phased, there was no need to phase it again.

**Supplementary Method 5. Minimac4 source code to obtain the θ value**

In Minimac4's implementation, the θ value was supplied by either providing it within the m3vcf file or using a reference genetic map.[5] The TOPMed imputation pipeline used the HapMap2 genetic map (see the main article). Minimac4 used linear interpolation to transform the recombination rate to the θ value between markers. By default, the transformed value is an internal variable. We modified the source code to output the values to the log file as follows:

(https://github.com/statgen/Minimac4/blob/v1.0.2/src/Analysis.cpp#L80)

```
for(i=0; i<referencePanel.numMarkers; i++)
{
     std::cout<<"Recom = "<<i<<"\t"<<referencePanel.Recom[i]<<std::endl;
}
```

#### Supplementary Notes

##### Supplementary Note 1. Consistency between MARE and $\beta_{imp}$ obtained from Rsq and dosage $r^2$ (or EmpRsq) and that calculated from imputed dosage and true genotype (or allele)

In calculating the MARE and $\beta_{imp}$ metrics, imputed dosage and true genotype (or true allele) were used. When haploid data was used, $\beta_{imp}$ was the regression slope between the imputed allelic dosage and true allele dose (encoded as 0 or 1), and MARE the residual sum of squares divided by $np(1-p)$ (the expected binomial variance on allele), where *n* is the number of haplotypes imputed, and *p* the alternative allele frequency (AAF) in imputed dataset. When diploid data was used, the linear regression was between the imputed dosage and true genotype (encoded as 0, 1, or 2), $\beta_{imp}$ was the regression slope, and MARE the residual sum of squares divided by $2np(1-p)$ (the expected variance on diploid genotype), where *n* is the number of samples imputed, and *p* the AAF.

As illustrated in the methods and results, three points may break the consistency between MARE and $\beta_{imp}$ calculated directly from the dosage and obtained from Rsq and dosage $r^2$ (or EmpRsq). They were as follows: (1) When dosage $r^2$ was calculated from diploid data (when external reference WGS was used) and Rsq from haploid data (Minimac4 reported Rsq in haploid form), Equations 6–7 would suffer from fluctuations caused by violating the Hardy-Weinberg equilibrium (HWE). (2) The AAF inconsistency between the imputed and WGS datasets affects the $\beta_{imp}$ in Equations 5 and 7. (3) Although Equations 6–7 did not contain AAF, Rsq and EmpRsq were not stable when the minor allele frequency (MAF) and minor allele count (MAC) were very low.[6] Here, we evaluated each point.

We categorized the variants into two groups according to the MAF of $WGS_{993}$: MAF ≥ 0.5% (MAC ≥ 10) and 0.5% > MAF > 0.1% (10 > MAC > 2). We then calculated the Pearson correlation between (1) AAF in the imputed dataset and $WGS_{993}$, (2) MARE calculated from the imputed dataset and obtained from Rsq and dosage $r^2$ (or EmpRsq), and (3) $\beta_{imp}$ calculated from the imputed dataset and obtained from Rsq and dosage $r^2$ (or EmpRsq).

The AAF was highly consistent for variants with MAF ≥ 0.5% ($r^2$ > 0.99) but only in modest or weak consistency for variants with 0.5% > MAF > 0.1% ($r^2$ = 0.07–0.56) (**Supplementary Figure 3**). The correlation of MARE was > 0.99 and > 0.94 for variants with MAF ≥ 0.5% and with 0.5% > MAF > 0.1% (**Supplementary Figure 4**). Calculating all metrics using the same haploid data fully diminished the fluctuations in MARE, as expected, because AAF discrepancy does not affect Equations 4 and 6 (**Supplementary Figure 5**). The correlation of $\beta_{imp}$ was > 0.91 and > 0.82 for variants with MAF ≥ 0.5% and with 0.5% > MAF > 0.1% (**Supplementary Figure 6**). It was also in good correlation (> 0.95 for variants with MAF ≥ 0.5%) when calculating from the same haploid data (**Supplementary Figure 7**).

These results showed that Equations 6–7 efficiently predict the estimated MARE and $\beta_{imp}$. Thus, on the Rsq ~ dosage $r^2$ (or EmpRsq) plot, each region corresponds to specific MARE and $\beta_{imp}$ values. We limited all downstream analyses to variants with a MAF ≥ 0.5% and MAC ≥ 10 to reduce fluctuations.

**Supplementary Note 2. Explanation of the θ value used in Minimac**

In Minimac's implementation, the θ value between the adjacent marker means there is a $P = 1 - \theta$ probability of not switching to other templates and a $P = \theta$ probability of randomly switching to other templates [5,7]. Thus, a low θ value between consecutive markers for a long genome region suggests long stretches of haplotype sharing [8]. The high spikes in the θ value suggest a high probability of recombination events [8]. Li et al. expected that the θ value reflects a combination of population recombination rates and the relatedness between the reference panel and the target samples (MaCH v1.0) [7]. In MaCH, both the samples in the reference panel and the target were used for estimating the θ value. Thus, in the current workflow of pre-phasing-imputation, the θ value only revealed the intrinsic characteristics of the reference panel [5]. In a typical imputation pipeline of Minimac3-Minimac4, the values are estimated from the reference panel using Minimac3 and stored in the m3vcf file. Here we scaled the θ value manually to study the impacts.

**Supplementary Note 3. Qualification and comparison of the θ value**

Because the θ value was between adjacent markers but not base pairs, the marker density would affect the θ estimates. It was infeasible to directly compare the θ value between large and small panels with different marker density. However, although the θ values between markers were different, the trend of total θ value along chromosome 19 did not change when constructing different reference panels using the same methods (**Supplementary Figure 14**), as expected, because recombination events were the intrinsic property of the reference panel. Hence, we used the total θ value for the comparison.

**Supplementary Note 4. Relationship between imputed-genotype certainty, Rsq, and INFO**

Let $x_i$ be the imputed allelic dosage, also the probability of alternative allele of the *i*-th haplotype, $X_i$. It has $P(X_i = 1|x_i) = x_i$ and $P(X_i = 0|x_i) = 1 - x_i$, which means the probability of $X_i$ being 1 is $x_i$. Given that $X_i$ only takes values 0 or 1, $X_i^2 = \begin{cases} 1, & if\ X_i = 1 \\ 0, & if\ X_i = 0 \end{cases}$, then:

$$E[X_i^2] = x_i \times 1 + (1 - x_i) \times 0 = x_i$$

(Equation 8)

$$E[X_i]^2 = x_i^2$$

(Equation 9)

Hence:

$$Var[X_i] = E[X_i^2] - E[X_i]^2 = x_i - x_i^2$$

(Equation 10)

$Var[X_i]$ is the variance of the *i*-th imputed allele. It is 0 when $x_i = 1 \text{ or } 0$, which means no uncertainty. And it is the maximum when $x_i = 0.5$, which means the true allele has the equal chance to be 0 or 1.

Given $X_i$ is independently imputed for each haplotype, the average variance for all alleles is:

$$\frac{\sum_{i=1}^{2N} Var[X_i]}{2N} = \frac{\sum_{i=1}^{2N}(x_i - x_i^2)}{2\text{N}}$$

(Equation 11)

where *2N* is the number of imputed haplotypes. Let $\boldsymbol{x} = (x_1, ..., x_{2N})$ a vector of the imputed allelic dosage and:

$$Var(\boldsymbol{x}) = \frac{\sum_{i=1}^{2N} x_i^2}{2N} - (\frac{\sum_{i=1}^{2N} x_i}{2N})^2 = \frac{\sum_{i=1}^{2N} x_i^2 - 2Np^2}{2N}$$

(Equation 12)

where $p = \frac{\sum_{i=1}^{2N} x_i}{2N}$ is the alternative allele frequency (AAF).

Add *2Np – 2Np* to the numerator of Equation 12:

$$Var(\boldsymbol{x}) = \frac{2Np - 2Np^2 - 2Np + \sum_{i=1}^{2N} x_i^2}{2N}$$

$$= \frac{2Np(1-p) - \sum_{i=1}^{2N} x_i + \sum_{i=1}^{2N} x_i^2}{2N}$$

$$= p(1-p) - \frac{\sum_{i=1}^{2N}(x_i - x_i^2)}{2N}$$

(Equation 13)

The first term in Equation 13 is the Binomial variance, given all alleles take values 0 or 1 (no uncertainty). The second term is shown in Equation 11. Because:

$$Rsq = \frac{Var(\boldsymbol{x})}{p(1-p)} = 1 - \frac{\sum_{i=1}^{2N}(x_i - x_i^2)}{2Np(1-p)}$$

(Equation 14)

the inverse relationship between Rsq and $Var[X_i]$ is thereby demonstrated. When all $x_i$ = *p*, there is no information gain from imputation, and Rsq = 0, as expected. When all $x_i$ take values 0 or 1, $x_i - x_i^2 = 0$, and Rsq = 1, also as expected.

Next, we demonstrate that Rsq equals INFO score. INFO is defined as [9]:

$$INFO = 1 - \frac{\sum_{\mathrm{j}=1}^{M}\left(4P_{jg2} + P_{jg1} - \left(x_{ja} + x_{jb}\right)^2\right)}{2Mp(1-p)}$$

(Equation 15)

Where *j* is the *j*-th individual and *ja jb* the imputed dosage of two alleles. $P_{jg1}$ and $P_{jg2}$ are probabilities of the imputed genotype being 1 or 2 for individual *j*. *M* is the number of individuals, and *p* the AAF. Given *ja* and *jb* are independent, $P_{jg1} = x_{ja}(1 - x_{jb}) + (1 - x_{ja})x_{jb}$ and $P_{jg2} = x_{ja}x_{jb}$. Then:

$$INFO = 1 - \frac{\sum_{j=1}^{M}(4x_{ja}x_{jb} + x_{ja}(1 - x_{jb}) + (1 - x_{ja})x_{jb} - (x_{ja} + x_{jb})^2)}{2Mp(1-p)}$$

$$= 1 - \frac{\sum_{j=1}^{M}(x_{ja} - x_{ja}^2 + x_{jb} - x_{jb}^2)}{2Mp(1-p)}$$

(Equation 16)

In the same dataset, $(x_{1a}, x_{1b}, \ldots, x_{Ma}, x_{Mb}) = (x_i, \ldots, x_{2N})$ and $M = N$, thus we have:

$$\sum\nolimits_{j=1}^{M}(x_{ja} - x_{ja}^2 + x_{jb} - x_{jb}^2) = \sum\nolimits_{i=1}^{2N}(x_i - x_i^2)$$

(Equation 17)

Finally, for a variant:

$$INFO = 1 - \frac{\sum_{i=1}^{2N}(x_i - x_i^2)}{2Np(1-p)} = Rsq$$

(Equation 18)

Taking Equations 11 and 18 together, Rsq and INFO are equal when calculating from the same dataset, and inversely associated with the average variance of imputed-allele. High certainty of an imputed allele decreases the variance and thereby increases Rsq and INFO.

**Supplementary Note 5. Explanation of the change in the θ value and the imputed allelic dosage**

We found that different θ values changed the imputed allelic dosage. The imputation is to find the unobserved paths through hidden states (corresponding to the reference haplotypes) to build an imperfect mosaic of templates that match the target haplotype while allowing allele mismatches [6]. The template switching rate (θ) is the probability that the path switches from one state to the next. The error rate (ε) represents the tolerance of mismatches between the selected mosaic path and the target haplotype. The posterior probability of the imputed allele is the sum of the state probabilities (for the missing markers, the emission probability of the state is 1) corresponding to reference haplotype(s) carrying that allele [6].

When running the imputation, only genotyped markers were available to find the paths. Switching and error rates could be interpreted as how easier a switch would happen and how many mismatches were acceptable. Given a specific ε value, switches are suppressed if all markers have θ values close to 0. The imputation algorithm would copy the most similar reference haplotype (*i.e.*, with the least mismatches) rather than finding a mosaic path. It would make the imputation result with less or no uncertainty (*i.e.*, all the imputed allelic dosages were equal to 0 or 1 as that of the selected reference haplotype) but with more wrongly imputed alleles [10], because this process does not always select haplotype(s) from the same ancestor (0.01–0.5 fold scaling of the θ value in **Supplementary Figure 12**). Conversely, if all markers have high θ values, many mosaic paths which could produce few errors in the genotyped markers were selected [10]. On average, it would shrink the imputed allelic dosage of missing markers to the reference panel's background AAF (2–100 fold scaling of the θ value in **Supplementary Figure 12**). If a marker has a higher ε value, then this marker is less weighted when computing the posterior probability of the path. In that case, all states obtain a more similar posterior probability, regardless of whether the allele is matched or mismatched between the template and target haplotype. Here we studied the θ value.

**Supplementary Note 6. Reference panel's impact on the θ estimates**

**Figure 6** conveys five crucial findings regarding the influence of reference panel composition on the θ estimates.

First, the total θ value decreased consistently (p = 7.85e-5 for each size increase, Wilcoxon rank-sum test, one-sided), with the trend being more modest as the JPT-only panel size increased (**Figure 6A**). This suggested that when the reference panel size was small (*e.g.*, < 2,000), adding samples made it easier to find shared haplotypes, hence expecting fewer switches. On the other hand, at a larger size, the plentiful haplotype sharing might make further addition of samples with modest effects.

Second, when the panel size was fixed and only change the ancestral composition, the total θ value decreased when mixing EUR with EAS (p=5.04e-4, Wilcoxon rank-sum test, one-sided) and then increased with further mixing AFR (p=7.85e-5, same as above), AMR (p=0.298, same as above), and SAS (p=7.85e-5, same as above) (**Figure 6B**). The ancestral diversity might make it harder to find shared haplotypes.

Third, in a EUR-EAS multi-ancestry panel, with the panel size increased from 403 to 4,163 and the proportion of EUR samples decreased from 100% to 9.68%, the total θ value decreased from 741.3 to 385.6, with a decrease of 52.0% (p=9.70e-4, 7.85e-5, 7.85e-5, 2.53e-4, 1.91e-4, 7.85e-5, 1.25e-5, for each size increase, Wilcoxon rank-sum test, one-sided) (**Figure 6C**). The ancestral diversity was more unbalanced in the large multi-ancestry reference panels. For example, the percentages of EAS samples in the HRC and TOPMed panels were 1.55% and 1.22%, respectively. Based on our simulation results, these reference panels might produce even lower θ value estimations than that estimated from an EAS-only reference panel with comparative sizes of the EAS samples in those large multi-ancestry panels.

Fourth, adding 1KGP-JPT to the $JPT_{3256}$ increased the total θ value slightly (an increment of 1.2%; p=0.013, Wilcoxon rank-sum test, one-sided) (**Figure 6D**). The total θ value increased with further addition of the remaining samples of 1KGP-EAS (p=1.43e-4, same as above), EUR (p=1.43e-4, same as above), AFR (p=1.91e-4, same as above), AMR (p=0.128, same as above), and SAS (p=6.31e-3, same as above). However, the change in the total θ value was from 327.7 to 443.8, with an increment of 35.4%, from $JPT_{3256}$ to $JPT_{3256}$+1KGP, revealing that even if the major ancestry group comprised only 65.28% of the samples in the reference panel, the change in the total θ value was relatively moderate.

Fifth, first with 104 1KGP-JPT, the total θ value decreased with the addition of the remaining EAS (p=7.85e-5, Wilcoxon rank-sum test, one-sided), EUR (p=7.85e-5, same as above), AFR (p=7.85e-5, same as above), and AMR (p=7.85e-5, same as above) (**Figure 6E**). It was revealed that the total θ value decreased as the sample size increased, regardless of the ancestry added, at the small panel size. Subsequently, the total θ value stopped decreasing at a sample size of approximately 2,000 with four ancestries. Further expanding to the five ancestries of 1KGP almost never changed the total θ value (p=0.910, Wilcoxon rank-sum test, two-sided). Combined with the simulation results from **Figures 6A–B**, the total θ value was in a trade-off between the panel size and ancestral diversity.

In summary, the θ estimates decreased with the sample size of a single ancestry and increased with ancestral diversity when size was fixed. While simultaneously increasing the panel size and ancestral diversity, the θ estimates were in a trade-off. The result also suggested that in a multi-ancestry panel of more than several thousand subjects, the θ estimates were underestimated for the minor ancestry and overestimated for the major ancestry compared to the value estimated in the single-ancestry panel.

#### Supplementary Note 7. Number of confident matches and high-Rsq variants obtained from distant ancestries using the 1KGP panel

We showed that no matter the target sample was the minor or major ancestry of the multi-ancestry reference panel, only a few variants existing on the haplotypes of distant ancestry could pass Rsq > 0.7. Thus, we made the conclusion that we would not expect to gain a significant number of high-Rsq variants (Rsq > 0.7) when combining the distant ancestry to expand the reference panel size. In scenarios 1 and 2 of the main article, we merged the JPT WGS and the 1KGP datasets by IMPUTE2. It has been reported that using IMPUTE2 to merge panels may bring variants from one panel to the missing sites of another panel [11]. Hence our methods may underestimate the number of well-imputed variants from each ancestry (in case a EUR variant was first imputed to the JPT reference haplotype and then to the target sample, it would neither be counted as EUR-only nor JPT-only).

To further quantify the contribution of high-Rsq variants from haplotypes of distant ancestry, we used the 1KGP panel. We defined the EAS-only and non-EAS variants in a straightforward manner. EAS-only variants only exist in the 504 EAS samples and non-EAS variants only exist in the 2,000 non-EAS samples. There were 83,023 EAS-only and 696,396 non-EAS variants. Using $WGS_{993}$ as the target sample, the number of confident matches (HDS > 0.9) was 98.9M (4.59% of the total variants × individual pairs), in which 185,870 or 3,362 belonged to EAS-only or non-EAS variants (the 1-fold θ value in **Supplementary Table 2**), respectively. Among the 159,469 high-Rsq variants (14.7% of the total variants), 8,627 and 724 were EAS-only or non-EAS, corresponding to a ratio of 10.4% and 0.10% of the total EAS-only and non-EAS variants, respectively (**Supplementary Table 2**). The result verified our conclusion that non-EAS variants were not rigorously presented in the imputation results.

#### Supplementary Figures

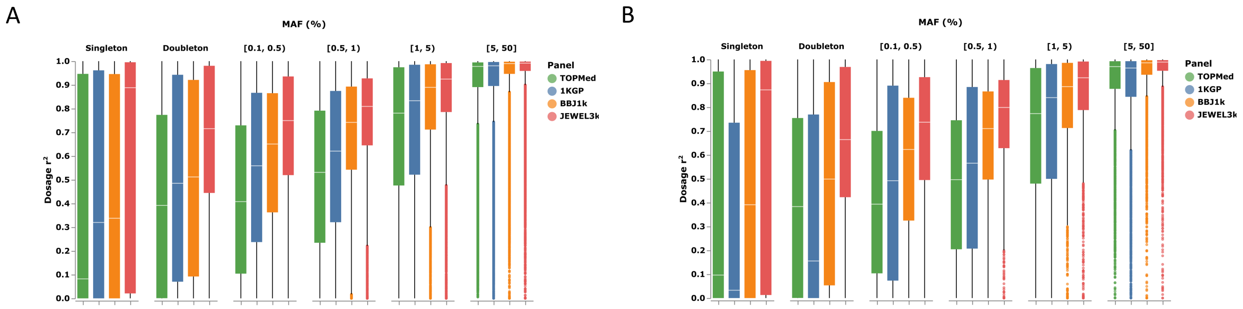


**Supplementary Figure 1. Dosage $r^2$ of all imputed variants.**

The box plot shows dosage $r^2$ of all imputed SNVs (A) and indels (B) (with Rsq ≥ 0.3 and in $WGS_{993}$) using the TOPMed, 1KGP, BBJ1k, and JEWEL3k reference panels, stratified by MAF of $WGS_{993}$. The boxes show the median, upper (75%), and lower (25%) quartiles. The whiskers show the 1.5-fold of the interquartile range (IQR) extended from the upper or lower quartile if there is a value exceeding them; otherwise, show the maximum or minimum value.

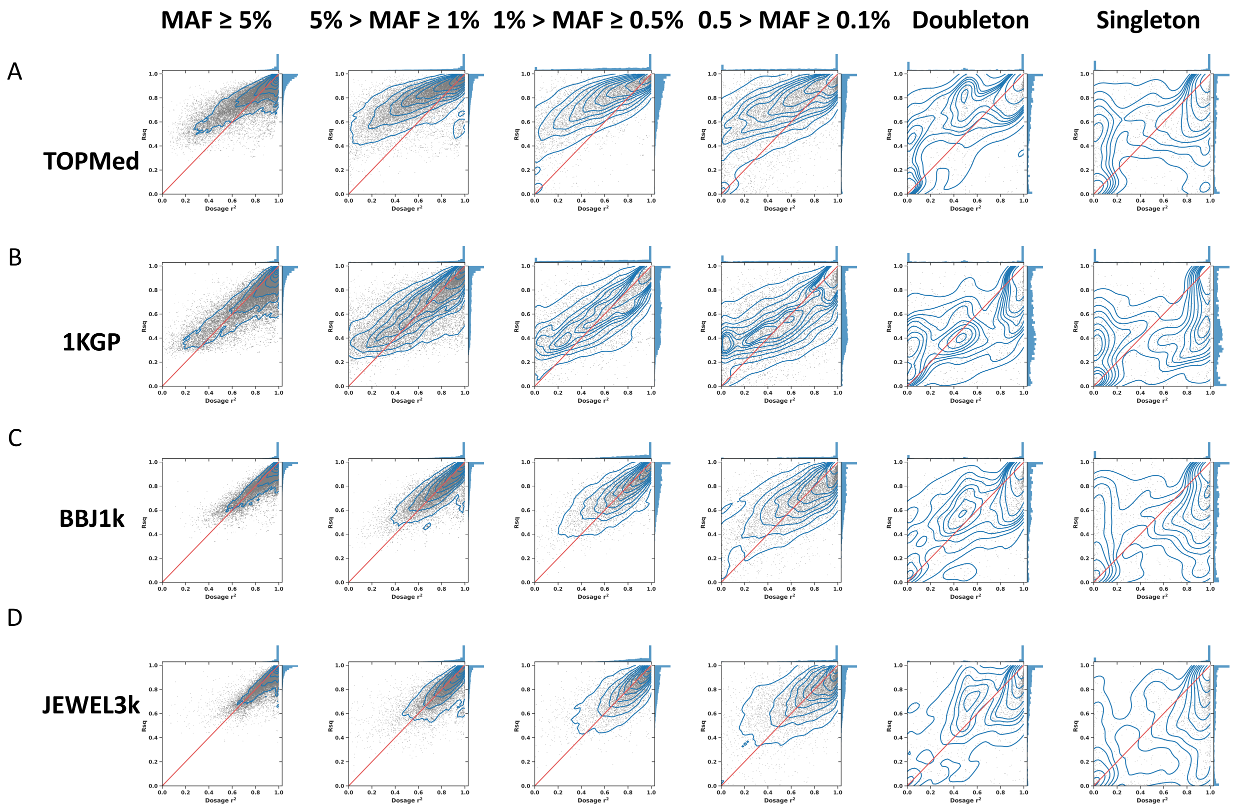


**Supplementary Figure 2. Comparison of dosage $r^2$ and Rsq in each MAF bin.**

The Supplementary Figure hows the deviation between dosage $r^2$ (x-axis) and Rsq (y-axis) in the TOPMed (A), 1KGP (B), BBJ1k (C), and JEWEL3k (D) imputation results. The scatter represents the variants, with the blue lines showing the density. The red line shows that Rsq equals dosage $r^2$, and the histograms on the side show the distribution. Overlapping variants imputed by the four panels were used. MAF is determined by $WGS_{993}$.

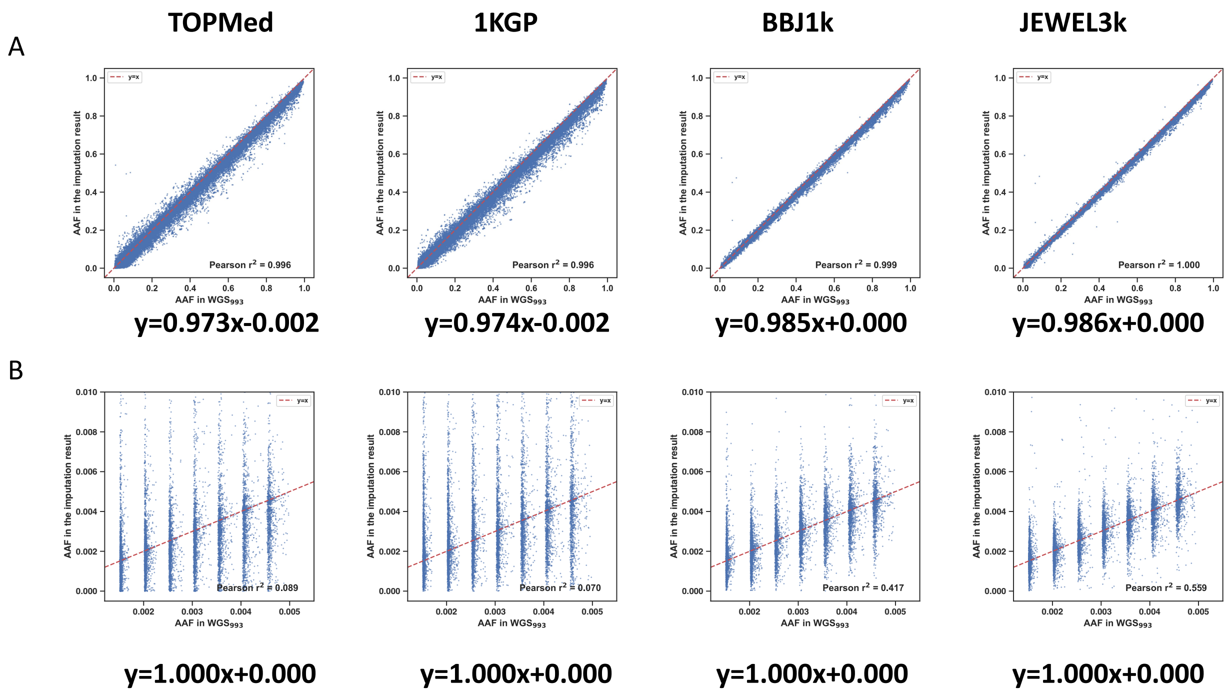


**Supplementary Figure 3. Comparison between the alternative allele frequency (AAF) of the imputed and WGS dataset.**

The scatter plot shows the comparison between AAF of the imputation result and $WGS_{993}$. Variants were grouped into (A) MAF ≥ 0.5% and (B) 0.5% > MAF ≥ 0.1% by MAF of $WGS_{993}$. In (B), variants with both 0.5% > MAF ≥ 0.1% and AAF < 0.5% are used.

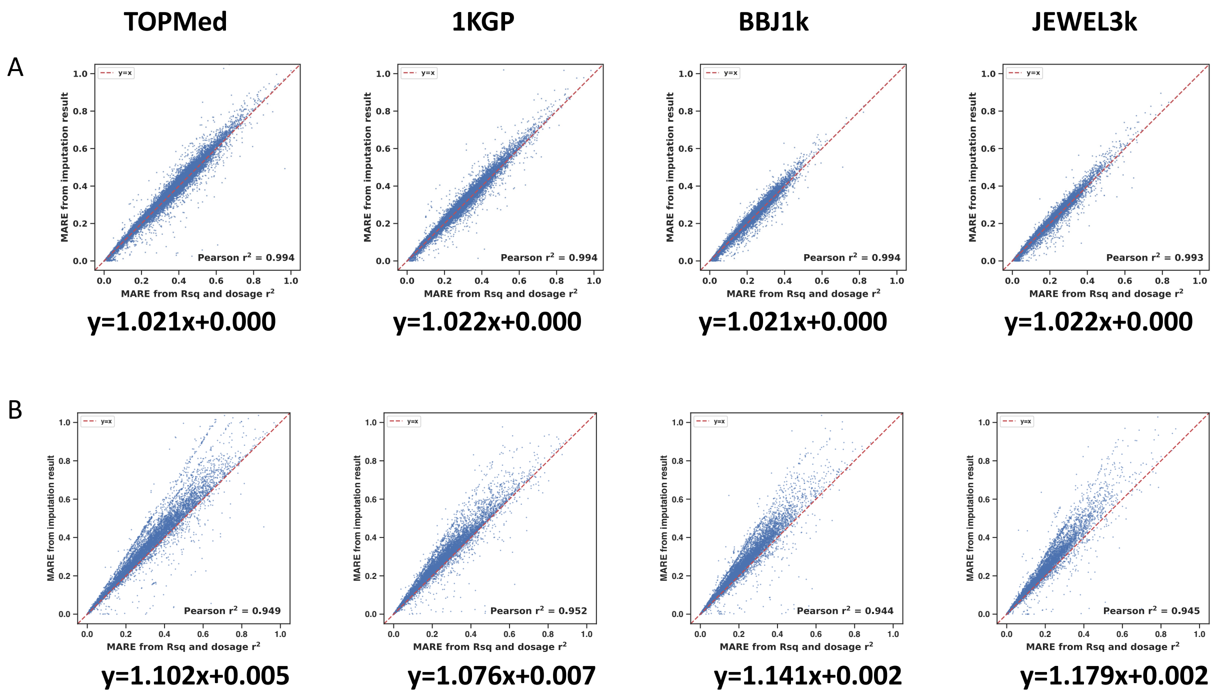


**Supplementary Figure 4. Comparison between MARE calculated from imputed dosage and obtained from Rsq and dosage $r^2$.**

The scatter plot shows the comparison between MARE calculated from imputation results or obtained from Rsq and dosage $r^2$. Variants were grouped into (A) MAF ≥ 0.5% and (B) 0.5% > MAF ≥ 0.1% by MAF of $WGS_{993}$.

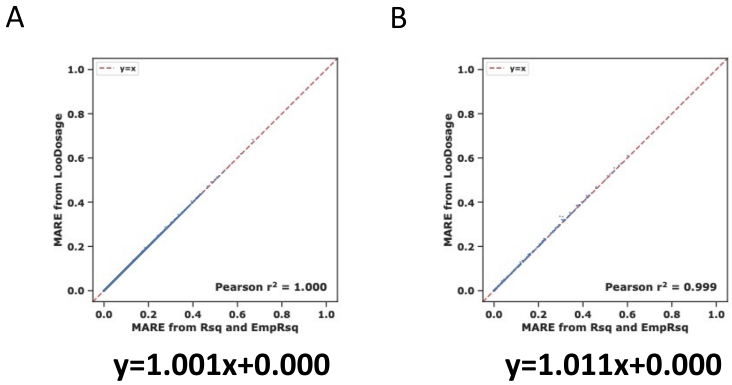


**Supplementary Figure 5. Comparison between MARE calculated from imputed allelic dosage and obtained from Rsq and EmpRsq.**

The scatter plot shows the comparison between MARE calculated from imputed allelic dosage (LooDosage) or obtained from Rsq and EmpRsq. LooDosage and EmpRsq are from the leave-one-out imputation of Minimac4 by hiding markers on the genotyping array. Variants were grouped into (A) MAF ≥ 0.5% and (B) 0.5% > MAF ≥ 0.1% by MAF of the genotyping array.

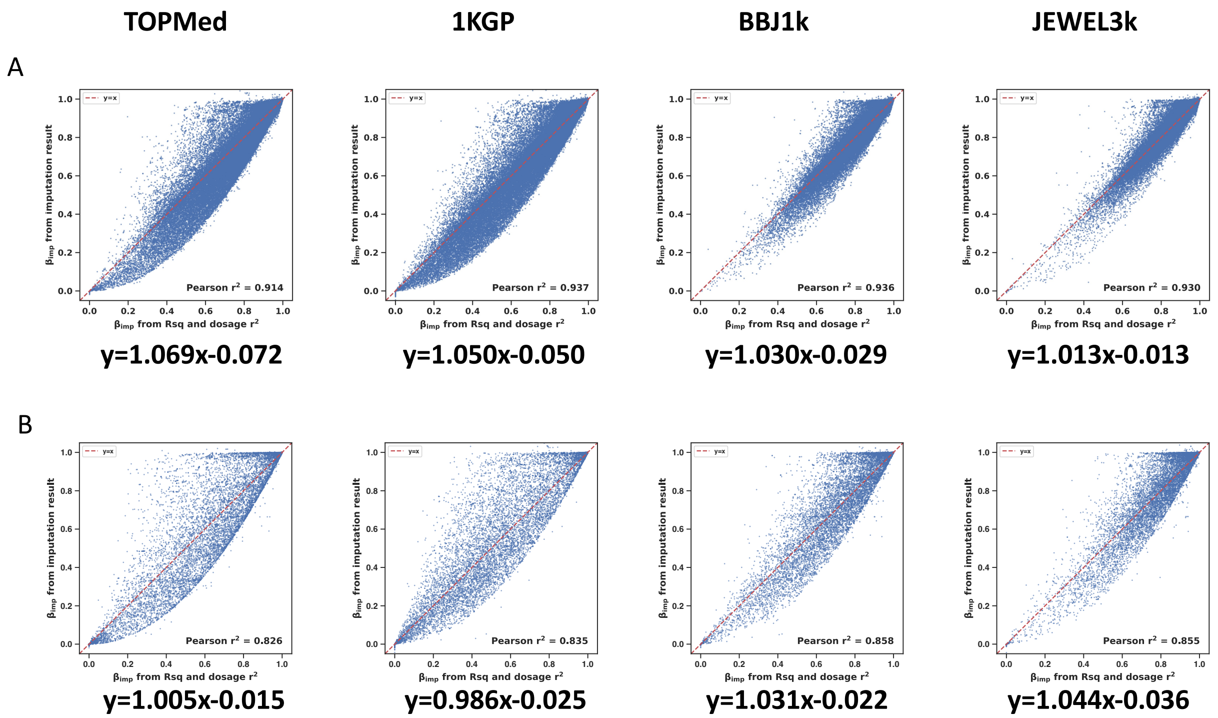


**Supplementary Figure 6. Comparison between $\beta_{imp}$ calculated from imputed dosage and obtained from Rsq and dosage $r^2$.**

The scatter plot shows the comparison between $\beta_{imp}$ calculated from imputation results or obtained from Rsq and dosage $r^2$. Variants were grouped into (A) MAF ≥ 0.5% and (B) 0.5% > MAF ≥ 0.1% by MAF of $WGS_{993}$.

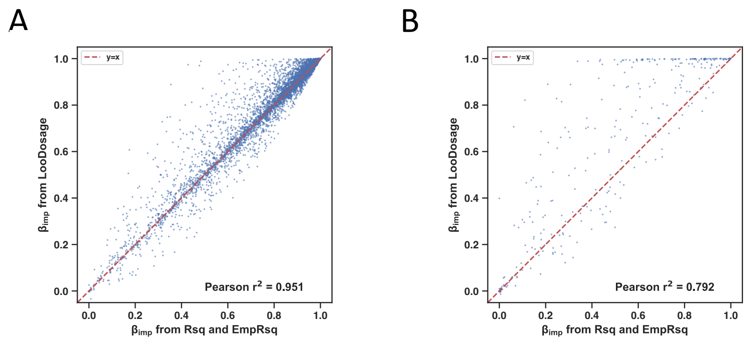


y=0.958x+0.043 y=1.016x+0.110

**Supplementary Figure 7. Comparison between $\beta_{imp}$ calculated from imputed allelic dosage and obtained from Rsq and EmpRsq.**

The scatter plot shows the comparison between $\beta_{imp}$ calculated from imputed allelic dosage (LooDosage) or obtained from Rsq and EmpRsq. Variants were grouped into (A) MAF ≥ 0.5% and (B) 0.5% > MAF ≥ 0.1% by MAF of the genotyping array.

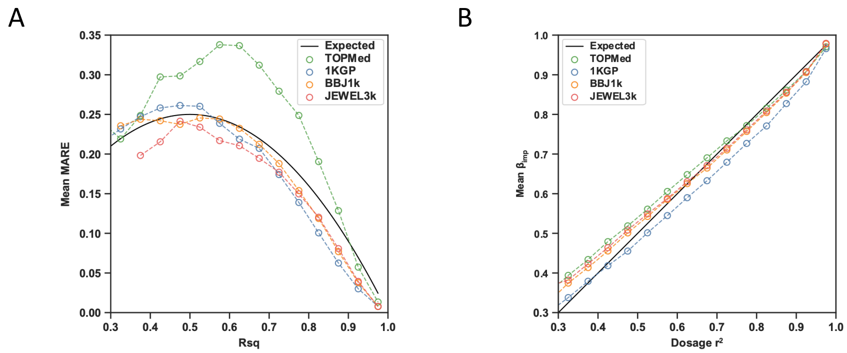


**Supplementary Figure 8. MARE and $\beta_{imp}$ of variants on chr19 using the TOPMed, 1KGP, BBJ1k, and JEWEL3k reference panels.**

(A) The mean MARE stratified by Rsq in each imputation result. (B) The mean $\beta_{imp}$ stratified by dosage $r^2$ in each imputation result. In (A) and (B), each bin has a width of 0.05, and bins with less than 50 variants are not shown. The expected values are calculated by assuming that Rsq equals dosage $r^2$. Overlapping variants imputed by the four panels were used. MAF is determined by $WGS_{993}$ and variants with MAF ≥ 0.5% were used.

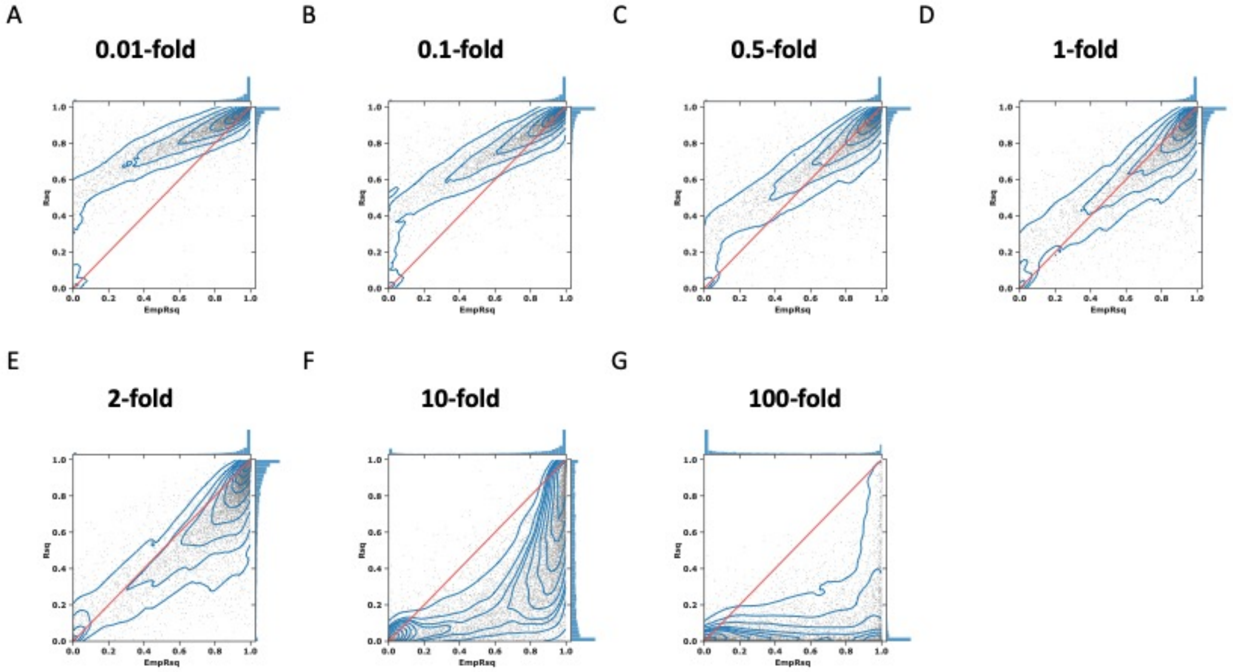


**Supplementary Figure 9. Comparison of EmpRsq and Rsq in the imputation results using different scalings of the θ value.**

The Supplementary Figure hows the deviation between EmpRsq (x-axis) and Rsq (y-axis) in the imputation results using the 1KGP reference panel, $WGS_{993}$ as the target sample, and seven scalings of the θ value. (A) 0.01-fold; (B) 0.1-fold; (C) 0.5-fold; (D) 1-fold (original); (E) 2-fold; (F) 10-fold; (G) 100-fold. The scatter represents the variants, with the blue lines showing the density. The red line shows that Rsq equals EmpRsq, and the histograms on the side show the distribution.

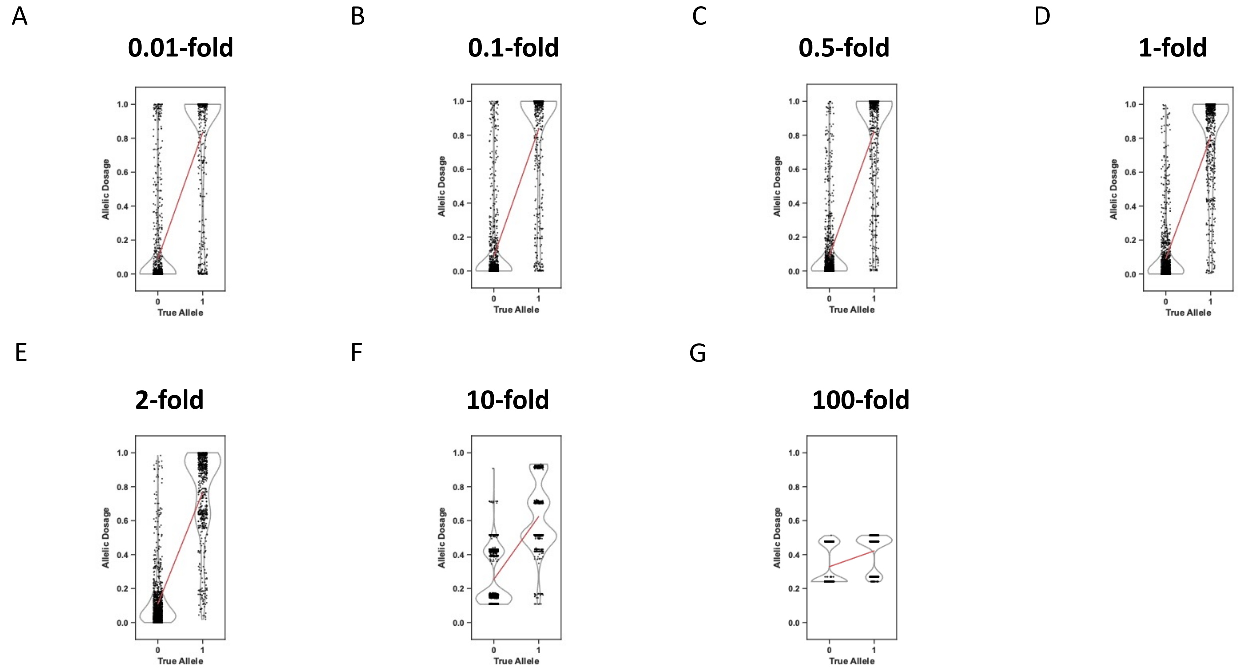


**Supplementary Figure 10. Allelic dosage distribution of rs1041062 in the imputation results using different scalings of the θ value.**

The Supplementary Figure hows the imputed allelic dosage of rs1041062 in the imputation results using the 1KGP reference panel, $WGS_{993}$ as the target sample, and seven scalings of the θ value. (A) 0.01-fold; (B) 0.1-fold; (C) 0.5-fold; (D) 1-fold (original); (E) 2-fold; (F) 10-fold; (G) 100-fold. The strip plots show the imputed allelic dosage and the violin plots show the distribution. The red line shows the linear regression line between the imputed allelic dosage and true allele.

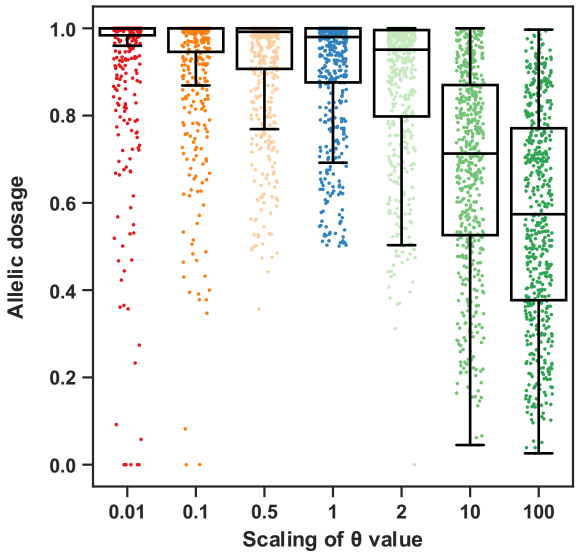


**Supplementary Figure 11. Distribution of the imputed allelic dosage with different scalings of the θ value.**

The Supplementary Figure hows the imputed allelic dosage of a randomly selected target haplotype (the same as Figure 4D) with the 7 scalings of the θ value. The scatters show variants. The boxes show the median, upper (75%), and lower (25%) quartiles. The whiskers show the 1.5-fold of the interquartile range (IQR) extended from the upper or lower quartile if there is a value exceeding them; otherwise, show the maximum or minimum value. Only variants with imputed allelic dosage > 0.5 and 0.3 < EmpRsq < 0.8 when using the 1-fold θ value are shown.

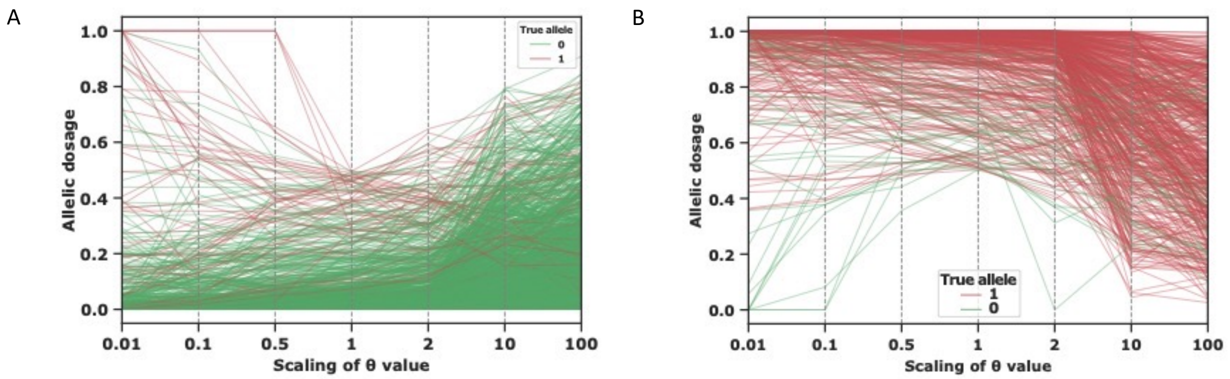


**Supplementary Figure 12. Changes in the imputed allelic dosage with different scalings of the θ value.**

The Supplementary Figure hows the changes in the imputed allelic dosage for the same target haplotype used in Figure 4D (A) and Supplementary Figure 11 (B). Each line represents a variant, the x-axis represents the scaling and y-axis represents the imputed allelic dosage. The two colors represent the true allele (0 or 1). A lower θ value only shrinks the dosages to 0 or 1 (increase certainty) but does not guarantee accuracy. On the other side, a higher θ value shrinks all imputed dosages to 0.5 (increase uncertainty).

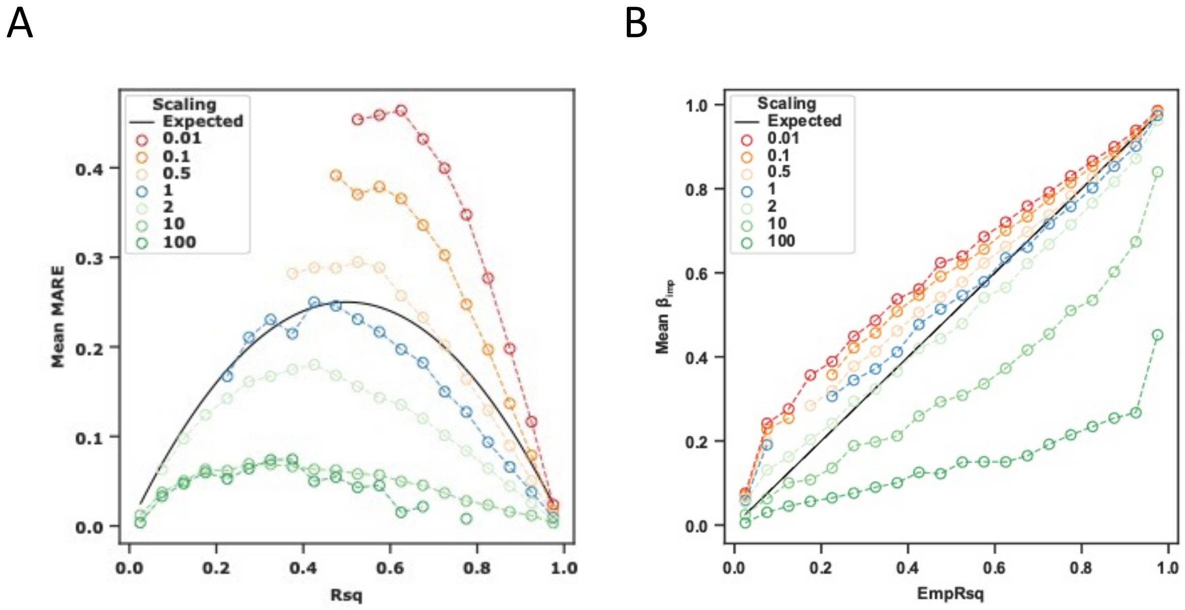


**Supplementary Figure 13. MARE and $\beta_{imp}$ of variants on chr19 using the 1KGP reference panel and different scalings of the θ value.**

(A) The mean MARE stratified by Rsq in each imputation result. (B) The mean $\beta_{imp}$ stratified by EmpRsq in each imputation result. In (A) and (B), each bin has a width of 0.05, and bins with less than 50 variants are not shown. The expected values are calculated by assuming that Rsq equals EmpRsq. MAF is determined by the genotyping array and variants with MAF ≥ 0.5% were used.

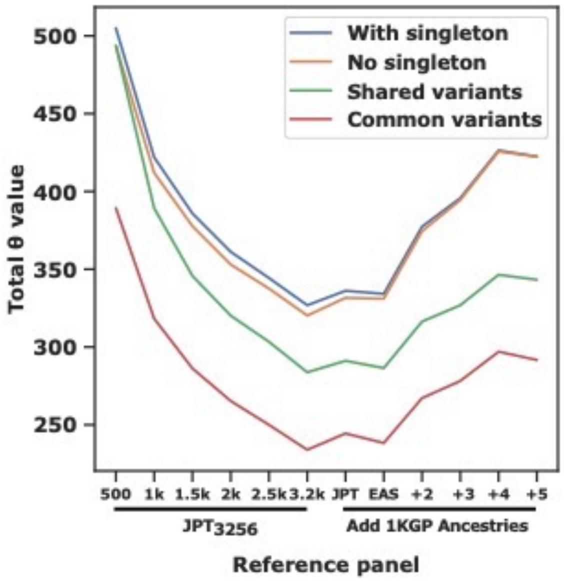


**Supplementary Figure 14. Total θ value along chromosome 19.**

The Supplementary Figure hows the total θ value along chromosome 19 estimated by Minimac3. Four different panel construction methods were compared. The x-axis labels are 500–3.2k: the JPT$_{3256}$ subset size; "JPT", EAS", and "+2"–"+5" ancestries: the 3,256 JPT + 1KGP-JPT, 1KGP-EAS, and the other 2–5 ancestries in the 1KGP, in the order of EUR, AFR, AMR, SAS. Panel construction methods are "With singleton": monomorphic variants and singletons are not removed when making subsets. These panels contain the same markers (identical number and position), with a size = 1.53M. "No singleton": monomorphic variants and singletons are removed. Number of markers in these panels varies from 510k to 1.53M. This is the routine strategy to build a reference panel for Minimac3/4. "Shared variants": all panels contain identical variants set, with a size of 370k (no singletons). "Common variants": each panel only contains variants with MAF ≥ 5%. MAF is determined by samples of each panel.

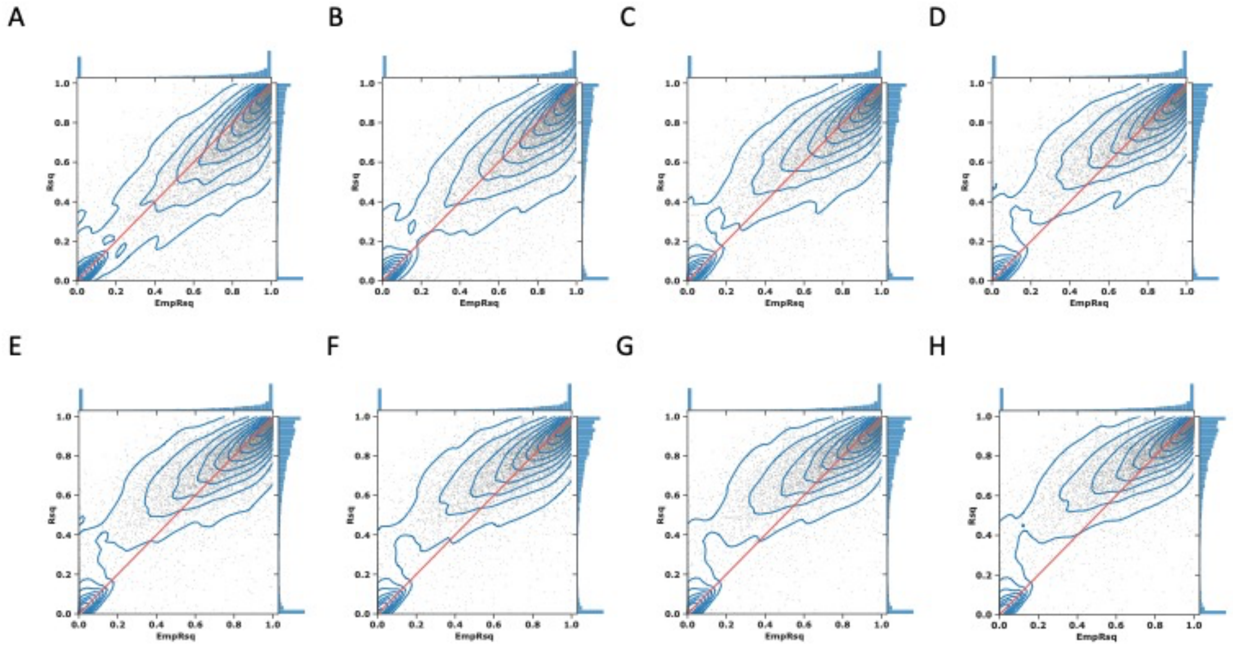


**Supplementary Figure 15. Comparison of EmpRsq and Rsq in the imputation results using simulated EUR-EAS reference panels.**

The Supplementary Figure hows the deviation between EmpRsq (x-axis) and Rsq (y-axis) in the imputation results using 8 simulated EUR-EAS reference panels. (A) EURn403; (B) EURn403 +1KGP-EAS; (C) EURn403 +1KGP-EAS+500JPT; (D) EURn403 +1KGP-EAS+1000JPT; (E) EURn403 +1KGP-EAS+1500JPT; (F) EURn403 +1KGP-EAS+2000JPT; (G) EURn403 +1KGP-EAS+2500JPT; (H) EURn403 +1KGP-EAS+3256JPT. The scatter represents the variants, with the blue lines showing the density. The red line shows that Rsq equals EmpRsq, and the histograms on the side show the distribution. EURn403 represents the 403 EUR; 500–3256JPT represents the number of JPT samples in the panel.

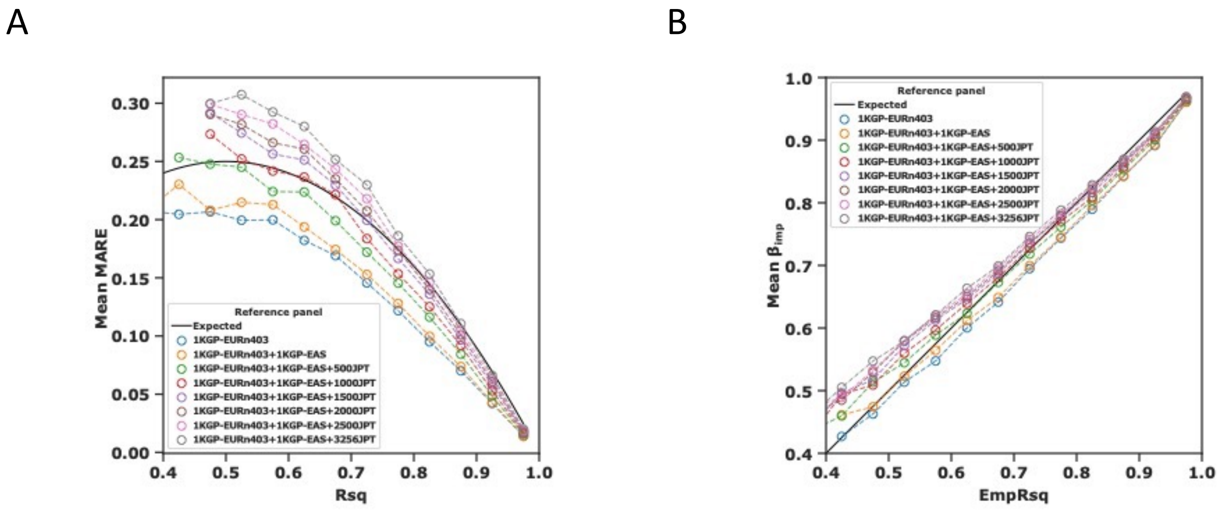


**Supplementary Figure 16. MARE and $\beta_{imp}$ of variants on chr19 using the simulated EUR-EAS reference panels.**

(A) The mean MARE stratified by Rsq in each imputation result. (B) The mean $\beta_{imp}$ stratified by EmpRsq in each imputation result. In (A) and (B), each bin has a width of 0.05, and bins with less than 50 variants are not shown. The expected values are calculated by assuming that Rsq equals EmpRsq. MAF is determined by the array and variants with MAF ≥ 0.5% were used. EURn403 represents the 403 EUR; 500–3256JPT represents the number of JPT samples in the panel.

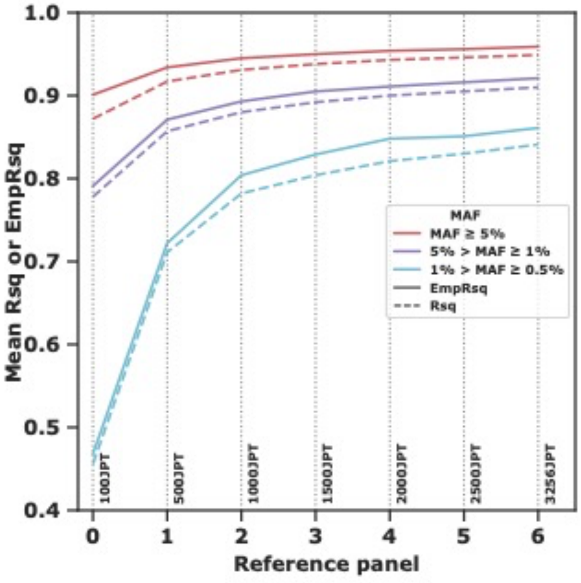


**Supplementary Figure 17. Mean EmpRsq and Rsq using JPT-only reference panels with different sizes.**

The plot shows the mean Rsq and EmpRsq of each imputation result, stratified by MAF of the array. The a-axis shows the reference panel indexes. Reference panels are indicated by the vertical dashed lines and labeled.

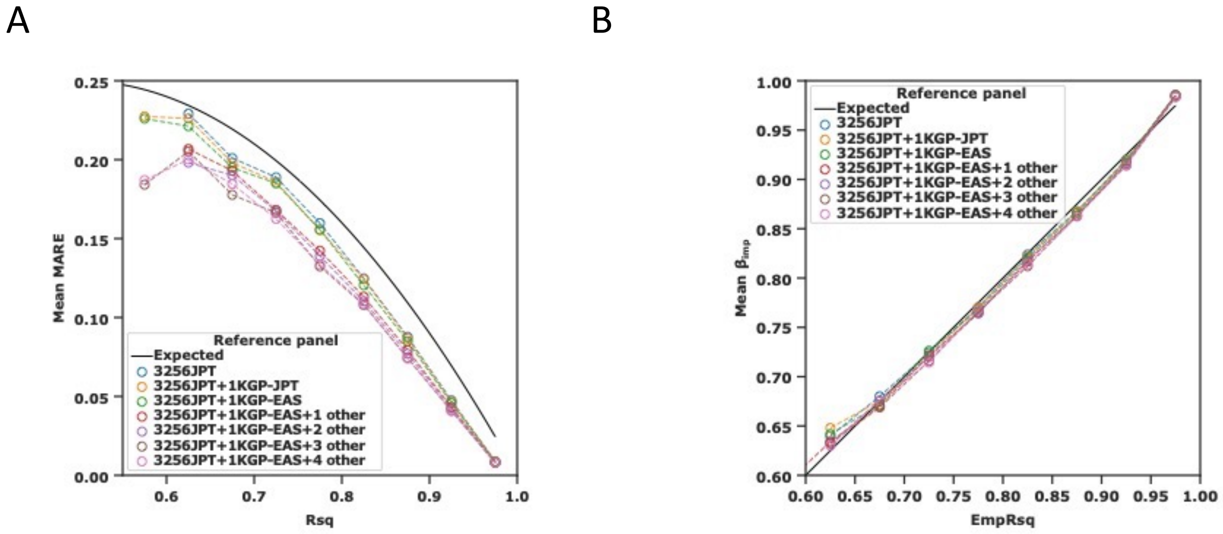


**Supplementary Figure 18. MARE and $\beta_{imp}$ of variants on hr19 using the 7 simulated JPT-1KGP reference panels.**

(A) The mean MARE stratified by Rsq in each imputation result. (B) The mean $\beta_{imp}$ stratified by EmpRsq in each imputation result. Each bin has a width of 0.05, and bins with less than 50 variants are not shown. The expected values are calculated by assuming that Rsq is equal to EmpRsq. MAF is determined by the array and variants with MAF ≥ 0.5% were used.

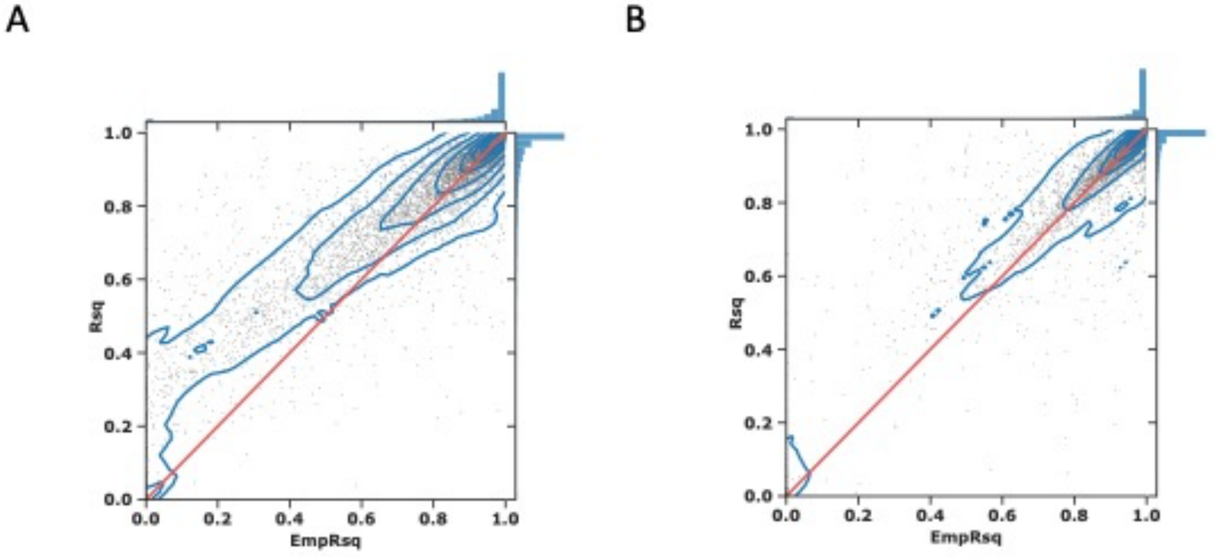


**Supplementary Figure 19. Comparison of the EmpRsq and Rsq in the imputation results using the HapMap2 genetic map as a reference for the θ value.**

The Supplementary Figure hows the deviation between EmpRsq and Rsq in the imputation results using the Minimac4 v1.0.2 and HapMap2 genetic map, the 1KGP (A) and JEWEL3k (B) as the reference panel, and $WGS_{993}$ as the target sample. The scatter represents the variants, with the blue lines showing the density. The red line shows that Rsq equals EmpRsq, and the histograms on the side show the distribution.

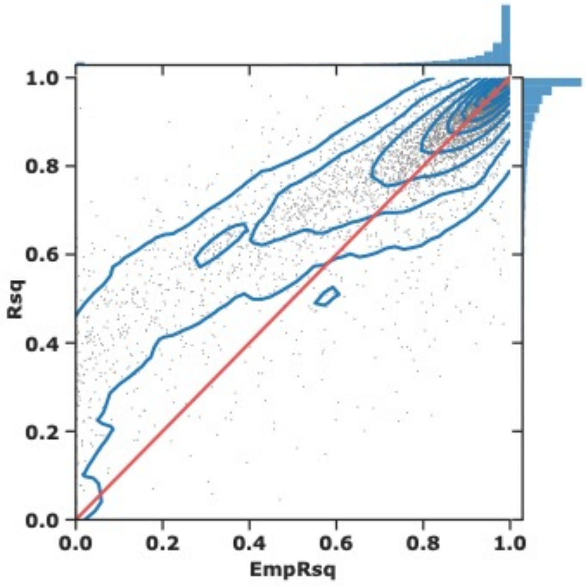


**Supplementary Figure 20. Comparison of the EmpRsq and Rsq in the imputation results using the HRC reference panel.**

The Supplementary Figure hows the deviation between EmpRsq and Rsq in the imputation results using the HRC reference panel (performed on the Michigan Imputation Server). The scatter represents the variants (on chr22), with the blue lines showing the density. The red line shows that Rsq equals EmpRsq, and the histograms on the side show the distribution.

#### Supplementary Tables

**Supplementary Table 1. Number of imputed variants using the four reference panels.**

| Reference panel | MAF (%) | SNV | | | Indel | | | All |
|---|---|---|---|---|---|---|---|---|
| | | Rsq ≥ 0 | Rsq ≥ 0.3 | Rsq ≥ 0.7 | Rsq ≥ 0 | Rsq ≥ 0.3 | Rsq ≥ 0.7 | Rsq ≥ 0 |
| TOPMed | [0, 0.05) | 252,771,622 | 7,573,456 | 1,218,495 | 19,491,389 | 553,114 | 88,286 | 272,263,011 |
| | [0.05, 0.5) | 8,592,816 | 5,424,023 | 2,182,279 | 681,658 | 434,383 | 169,287 | 9,274,474 |
| | [0.5, 1) | 1,313,057 | 1,075,612 | 738,434 | 104,566 | 87,431 | 58,194 | 1,417,623 |
| | [1, 5) | 2,324,580 | 1,979,385 | 1,633,050 | 176,655 | 156,127 | 126,582 | 2,501,235 |
| | [5, 50] | 5,387,174 | 5,339,524 | 5,150,778 | 370,890 | 368,148 | 354,359 | 5,758,064 |
| | All | 270,389,249 | 21,392,000 | 10,923,036 | 20,825,158 | 1,599,203 | 796,708 | 291,214,407 |
| 1KGP | [0, 0.05) | 26,375,615 | 913,226 | 185,101 | 1,484,317 | 60,437 | 13,708 | 27,859,932 |
| | [0.05, 0.5) | 8,201,495 | 2,075,814 | 502,547 | 601,000 | 149,797 | 34,227 | 8,802,495 |
| | [0.5, 1) | 1,262,622 | 803,635 | 357,303 | 112,610 | 70,492 | 26,261 | 1,375,232 |
| | [1, 5) | 2,318,671 | 1,825,146 | 1,251,700 | 249,258 | 188,133 | 116,457 | 2,567,929 |
| | [5, 50] | 5,646,789 | 5,518,738 | 5,141,588 | 857,088 | 832,699 | 690,649 | 6,503,877 |
| | All | 43,805,192 | 11,136,559 | 7,438,239 | 3,304,273 | 1,301,558 | 881,302 | 47,109,465 |
| BBJ1k | [0, 0.05) | 33,475,711 | 5,351,085 | 959,548 | 1,601,355 | 243,654 | 47,694 | 35,077,066 |
| | [0.05, 0.5) | 13,816,241 | 10,986,526 | 3,330,407 | 758,207 | 592,295 | 178,565 | 14,574,448 |
| | [0.5, 1) | 1,497,074 | 1,303,683 | 801,332 | 112,479 | 95,378 | 55,523 | 1,609,553 |
| | [1, 5) | 2,253,326 | 2,059,123 | 1,691,900 | 182,249 | 162,828 | 129,736 | 2,435,575 |
| | [5, 50] | 5,258,811 | 5,212,583 | 5,085,955 | 431,617 | 425,821 | 402,839 | 5,690,428 |
| | All | 56,301,163 | 24,913,000 | 11,869,142 | 3,085,907 | 1,519,976 | 814,357 | 59,387,070 |
| JEWEL3k | [0, 0.05) | 44,913,596 | 14,103,823 | 3,481,903 | 2,111,008 | 680,368 | 181,987 | 47,024,604 |
| | [0.05, 0.5) | 11,895,969 | 10,686,265 | 4,654,377 | 661,730 | 595,772 | 276,365 | 12,557,699 |
| | [0.5, 1) | 1,374,793 | 1,206,608 | 866,849 | 95,832 | 81,573 | 58,256 | 1,470,625 |
| | [1, 5) | 2,194,888 | 2,041,329 | 1,775,366 | 162,166 | 148,384 | 127,112 | 2,357,054 |
| | [5, 50] | 5,279,010 | 5,224,012 | 5,111,624 | 387,791 | 383,145 | 373,344 | 5,666,801 |
| | All | 65,658,256 | 33,262,037 | 15,890,119 | 3,418,527 | 1,889,242 | 1,017,064 | 69,076,783 |

Minor allele frequency (MAF) is determined by each imputed dataset.

**Supplementary Table 2. Number of confident alleles and high-Rsq variants using the 1KGP reference panel and 21 scalings of the θ value.**

| Scaling of θ value | Alleles with HDS > 0.9 | | | Variants with Rsq > 0.7 | | |
|---|---|---|---|---|---|---|
| | All | EAS-only | Non-EAS | All | EAS-only | Non-EAS |
| 0.01 | 111,900,629 | 353,418 | 41,140 | 214,832 | 20,923 | 7,213 |
| 0.02 | 111,419,293 | 340,835 | 34,699 | 209,018 | 19,440 | 5,858 |
| 0.1 | 109,378,239 | 303,045 | 19,783 | 194,113 | 15,882 | 2,999 |
| 0.125 | 108,895,828 | 295,656 | 18,003 | 191,782 | 15,301 | 2,665 |
| 0.2 | 107,693,098 | 278,442 | 14,059 | 186,726 | 14,019 | 2,072 |
| 0.25 | 107,009,589 | 268,947 | 12,108 | 184,184 | 13,428 | 1,864 |
| 0.33 | 106,046,221 | 255,937 | 9,914 | 180,399 | 12,606 | 1,569 |
| 0.5 | 104,270,414 | 233,269 | 7,111 | 173,640 | 11,198 | 1,218 |
| 0.67 | 102,479,229 | 215,508 | 5,251 | 168,236 | 10,220 | 936 |
| 0.8 | 101,026,131 | 203,389 | 4,356 | 164,512 | 9,544 | 819 |
| 1 | 98,875,870 | 185,870 | 3,362 | 159,469 | 8,627 | 724 |
| 1.25 | 95,910,630 | 166,420 | 2,636 | 153,657 | 7,626 | 608 |
| 1.5 | 92,957,791 | 148,619 | 2,031 | 147,868 | 6,821 | 490 |
| 2 | 87,221,001 | 118,354 | 1,325 | 137,454 | 5,400 | 371 |
| 3 | 75,388,680 | 70,260 | 785 | 118,479 | 3,210 | 264 |
| 4 | 64,177,890 | 38,507 | 662 | 101,666 | 1,826 | 203 |
| 5 | 54,572,785 | 22,061 | 567 | 88,379 | 1,118 | 175 |
| 8 | 38,237,092 | 10,462 | 445 | 65,357 | 594 | 147 |
| 10 | 32,070,626 | 9,027 | 469 | 56,827 | 525 | 140 |
| 50 | 17,265,686 | 7,431 | 456 | 30,874 | 391 | 119 |
| 100 | 16,297,937 | 7,423 | 443 | 28,538 | 387 | 117 |

The number of EAS-only and non-EAS variants were 83,023 and 696,396.

**Supplementary Table 3. Number of variants, confident alleles, and high-Rsq variants using using the simulated EUR-EAS reference panels with different sizes and EUR proportions, and 100 EUR as the target sample.**

| Reference panel | Number of variants in reference panel | | Alleles with HDS > 0.9 | | | Variants with Rsq > 0.7 | | |
|---|---|---|---|---|---|---|---|---|
| | All | Non-EUR | All | EUR-only | Non-EUR | All | EUR-only | Non-EUR |
| 1KGP-EURn403 | 456,835 | * | 8,670,087 | 84,797 | * | 163,468 | 18,646 | * |
| 1KGP-EURn403+1KGP-EAS | 815,407 | 358,572 | 8,523,623 | 85,099 | 181 | 168,046 | 20,811 | 242 |
| 1KGP-EURn403+1KGP-EAS+500JPT | 912,663 | 455,828 | 8,697,251 | 90,731 | 260 | 179,340 | 23,221 | 296 |
| 1KGP-EURn403+1KGP-EAS+1000JPT | 978,746 | 521,911 | 8,787,978 | 93,271 | 284 | 184,613 | 24,472 | 308 |
| 1KGP-EURn403+1KGP-EAS+1500JPT | 1,030,657 | 573,822 | 8,901,041 | 96,251 | 337 | 190,618 | 25,731 | 395 |
| 1KGP-EURn403+1KGP-EAS+2000JPT | 1,073,079 | 616,244 | 8,930,669 | 97,674 | 363 | 193,091 | 26,392 | 432 |
| 1KGP-EURn403+1KGP-EAS+2500JPT | 1,108,702 | 651,867 | 8,969,106 | 99,053 | 405 | 195,843 | 27,024 | 458 |
| 1KGP-EURn403+1KGP-EAS+3256JPT | 1,152,852 | 696,017 | 9,039,186 | 101,116 | 458 | 200,161 | 27,941 | 521 |

The number of EUR-only variants was 114,606. (*) denotes not available. EURn403 represents the 403 EUR; 1KGP-EAS represents the 504 EAS; 500–3256JPT represents the number of JPT samples in the reference panel.

**Supplementary Table 4. Number of variants, confident alleles, and high-Rsq variants using the simulated JPT-1KGP reference panels with different sizes and ancestral diversities, and $WGS_{993}$ as the target sample.**

| Reference panel | Number of variants in reference panel | | Alleles with HDS > 0.9 | | | | Variants with Rsq > 0.7 | | | |
|---|---|---|---|---|---|---|---|---|---|---|
| | All | Non-EAS | All | $JPT_{3256}$-only | 1KGP-EAS-only | Non-EAS | All | $JPT_{3256}$-only | 1KGP-EAS-only | Non-EAS |
| 100JPT | 295,966 | * | * | * | * | * | 136,682 | * | * | * |
| 500JPT | 510,113 | * | * | * | * | * | 178,534 | * | * | * |
| 1000JPT | 662,071 | * | * | * | * | * | 210,849 | * | * | * |
| 1500JPT | 769,906 | * | * | * | * | * | 232,582 | * | * | * |
| 2000JPT | 855,094 | * | * | * | * | * | 249,553 | * | * | * |
| 2500JPT | 924,533 | * | * | * | * | * | 261,409 | * | * | * |
| 3256JPT | 1,010,230 | * | 96,451,515 | 24,599 | * | * | 274,343 | 10,613 | * | * |
| 3256JPT+1KGP-JPT | 1,011,682 | * | 96,286,919 | 24,171 | * | * | 272,521 | 10,554 | * | * |
| 3256JPT+1KGP-EAS | 1,038,246 | * | 96,190,813 | 24,151 | 159 | * | 271,937 | 10,884 | 158 | * |
| 3256JPT+1KGP-EAS+EUR | 1,163,109 | 124,863 | 95,590,037 | 23,488 | 138 | 183 | 266,300 | 10,692 | 144 | 184 |
| 3256JPT+1KGP-EAS+EUR+AFR | 1,471,007 | 432,761 | 95,062,865 | 23,891 | 143 | 200 | 267,008 | 10,893 | 143 | 219 |
| 3256JPT+1KGP-EAS+EUR+AFR+AMR | 1,486,412 | 448,166 | 94,503,390 | 23,383 | 130 | 229 | 263,233 | 10,775 | 137 | 201 |
| 3256JPT+1KGP-All | 1,533,735 | 495,489 | 94,488,850 | 23,666 | 131 | 254 | 264,221 | 11,015 | 142 | 265 |

The number of $JPT_{3256}$-only and 1KGP-EAS-only variants was 74,490 and 7,627. (*) denotes not available. 100–3256 JPT represents the number of JPT samples in the reference panel; 1KGP- followed by the ancestry represents the 1KGP subset.

**Supplementary Table 5. The total θ value estimated by Minimac3 or transformed from the HapMap2 genetic map using Minimac4 v1.0.2.**

| Reference panel | Parameter source | Background template switching rate | Number of variants with a background template switching rate | Number of variants not with the background template switching rate | Total θ value | Ratio between the transformed and estimated total θ values |
|---|---|---|---|---|---|---|
| 1KGP | Genetic map and Minimac4 | 1.00E-05 | 493,335 | 591,200 | 110.4 | 0.152 |
| JEWEL3k | Genetic map and Minimac4 | 1.00E-05 | 792,501 | 741,234 | 112.6 | 0.267 |
| 1KGP | Minimac3 | 0.00065347 | 1,084,368 | 166 | 728 | * |
| JEWEL3k | Minimac3 | 0.00027504 | 1,533,730 | 4 | 422.4 | * |

Background template switching rate is a minimum probability of switching Minimac sets for all markers. (*) denotes not available.
